# Coordinated circadian timing through the integration of local inputs in *Arabidopsis thaliana*

**DOI:** 10.1101/617803

**Authors:** Mark Greenwood, Mirela Domijan, Peter D. Gould, Anthony J.W. Hall, James C.W. Locke

## Abstract

Every plant cell has a genetic circuit, the circadian clock, that times key processes to the day-night cycle. These clocks are aligned to the day-night cycle by multiple environmental signals that vary across the plant. How does the plant integrate clock rhythms, both within and between organs, to ensure coordinated timing? To address this question, we examined the clock at the sub-tissue level across *Arabidopsis thaliana* seedlings under multiple environmental conditions and genetic backgrounds. Our results show that the clock runs at different speeds (periods) in each organ, which causes the clock to peak at different times across the plant in both constant environmental conditions and light-dark cycles. Closer examination reveals that spatial waves of clock gene expression propagate both within and between organs. Using a combination of modeling and experiment, we reveal that these spatial waves are the result of the period differences between organs and local coupling, rather than long distance signaling. With further experiments we show that the endogenous period differences, and thus the spatial waves, are caused by the organ specificity of inputs into the clock. We demonstrate this by modulating periods using light and metabolic signals, as well as with genetic perturbations. Our results reveal that plant clocks are set locally by organ specific inputs, but coordinated globally via spatial waves of clock gene expression.

## Introduction

In response to the Earth’s predictable light-dark (LD) cycles, many organisms have evolved a circadian clock (1). A common design principle is a central oscillator that receives input from multiple environmental signals and uses them to predict the time of day. This timing information is used to coordinate processes, matching them to the optimum time of day or year. In plants, these processes include photosynthesis, leaf movement, and flowering (2).

A number of studies have reported that different parts of the plant can generate circadian oscillations with different periods under constant conditions (3). This could be due to the clock network being wired differently in different parts of the plant, or that the sensitivity of the clock to environmental inputs varies across the plant. There is already some evidence that both the network and inputs have some cell or tissue specificity. Previous work has shown that although most clock genes are expressed in most cell types (4–7), some core clock genes have a tissue enriched expression pattern (4,8,9). Additionally, it has been shown that different cell types respond preferentially to temperature or light inputs (10–12), and that the shoot and root clock have different sensitivities to light (6). However, how whole-plant timing is affected by tissue level differences in the clock network, or differences in sensitivity to clock inputs, remains unclear. In complex organisms, many physiological processes, including those under control of the clock, require coordinated timing across tissues. In many eukaryotes, cell-cell communication maintains clock coherence across the organism. For example, in mammals clock cells located in the suprachiasmatic nucleus drive rhythms across the body via neural and humoral signals (1,13). In plants, studies of synchronization (5,14–19), grafting experiments (18), and the use of tissue specific promoters (20) suggest cell-cell communication is also important for coherent rhythms. It has been proposed that this communication acts hierarchically, with the root clock dependent on a long-distance signal from the shoot (9,18,21). However, a decentralized structure, with multiple points of coordination across the plant, could potentially explain inconsistencies such as fast cells in the root tip (5), spiral and striped expression patterns in leaves and roots (14–16,22–24), and the entrainment of detached roots by light (6,25). Therefore, how plants coordinate the clock at the organism level is not understood (Fig 1). More specifically, it is not known whether cell-cell communication acts through local or long-distance signaling pathways.

**Fig 1.**
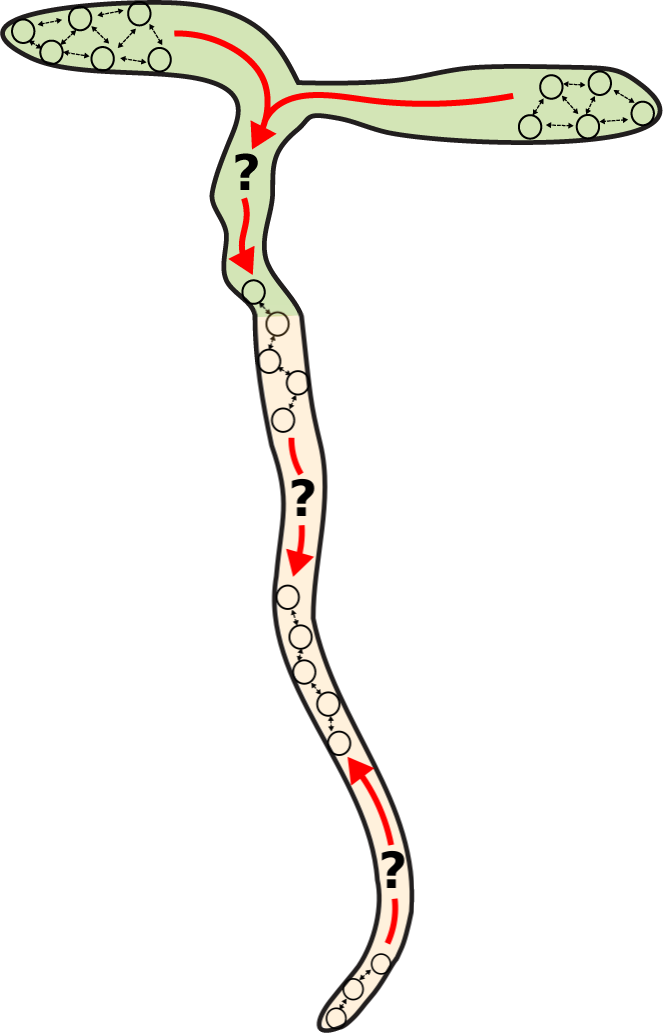
How do circadian clocks in different organs coordinate together? Individual clocks could communicate both within (black arrows) and between (red arrows) organs in order to coordinate plant timing.

In this work, we examined the clock at the sub-tissue level across *A. thaliana* seedlings *in vivo*. We observed that each organ of the plant has a different clock phase, even under LD cycles. Sub-tissue level analysis revealed that spatial waves of clock gene expression propagate within and between the organs. Mathematical models propose that waves under both constant light and LD cycles could be due to the combination of different periods in each part of the plant and local cell-cell coupling. We tested these predictions by examining rhythms in dissected plant roots. Waves up and down the root persisted in detached roots, showing that long distance signals from the shoot are not required for coordination. Next, by modulating periods in specific parts of the plant using genetic and environmental perturbations, we found that we could alter wave generation in a predictable manner. Thus, the clock in plants has a decentralized structure, with clocks across the plant coordinating via local cell-cell signaling.

## Results

### Organ specific clocks entrain to LD cycles with different phases

To investigate the coordination of clock rhythms, we analyzed rhythms across entire seedlings under different entrainment regimes. To do this, we monitored promoter activity of the core clock gene *GIGANTEA* (*GI*; 22) fused to the *LUCIFERASE* (*LUC*) reporter gene, for multiple days at near cellular resolution (Materials and Methods). This reporter line was chosen due to its strong expression, and its similar spatial expression to other clock components (5).

In order to observe the endogenous component of the rhythms, we first imaged seedlings under constant light (LL), having previously grown them under LD cycles (LD-to-LL; Fig 2A and Materials and Methods). Under the LD-to-LL condition we observed phase differences of *GI*::*LUC* expression between organs (Fig 2B, C). The cotyledon and hypocotyl peaked before the root, but the tip of the root peaked before the middle region of the root (Fig 2C, S1 Fig and S1 Video). Further, we observed a decrease in coherence between regions over time, with a range between the earliest and latest peaking region of 4.92 ± 3.79 h in the first and 18.36 ± 5.67 h in the final oscillation. This is due to the emergence of period differences between all regions (Fig 2D). The cotyledon maintained a mean period of 23.82 ± 0.60 h, whereas the hypocotyl and root ran at 25.41 ± 0.91 h and 28.04 ± 0.86 h respectively. However, the root tip ran slightly faster than the middle of the root, with a mean period of 26.90 ± 0.45 h, demonstrating the presence of endogenous period differences across all regions. These observations are qualitatively similar to the periods and phases previously observed in isolated organs (6,18,21), and at the cellular level across the seedling (5), validating our whole-plant assay for the circadian clock.

**Fig 2.**
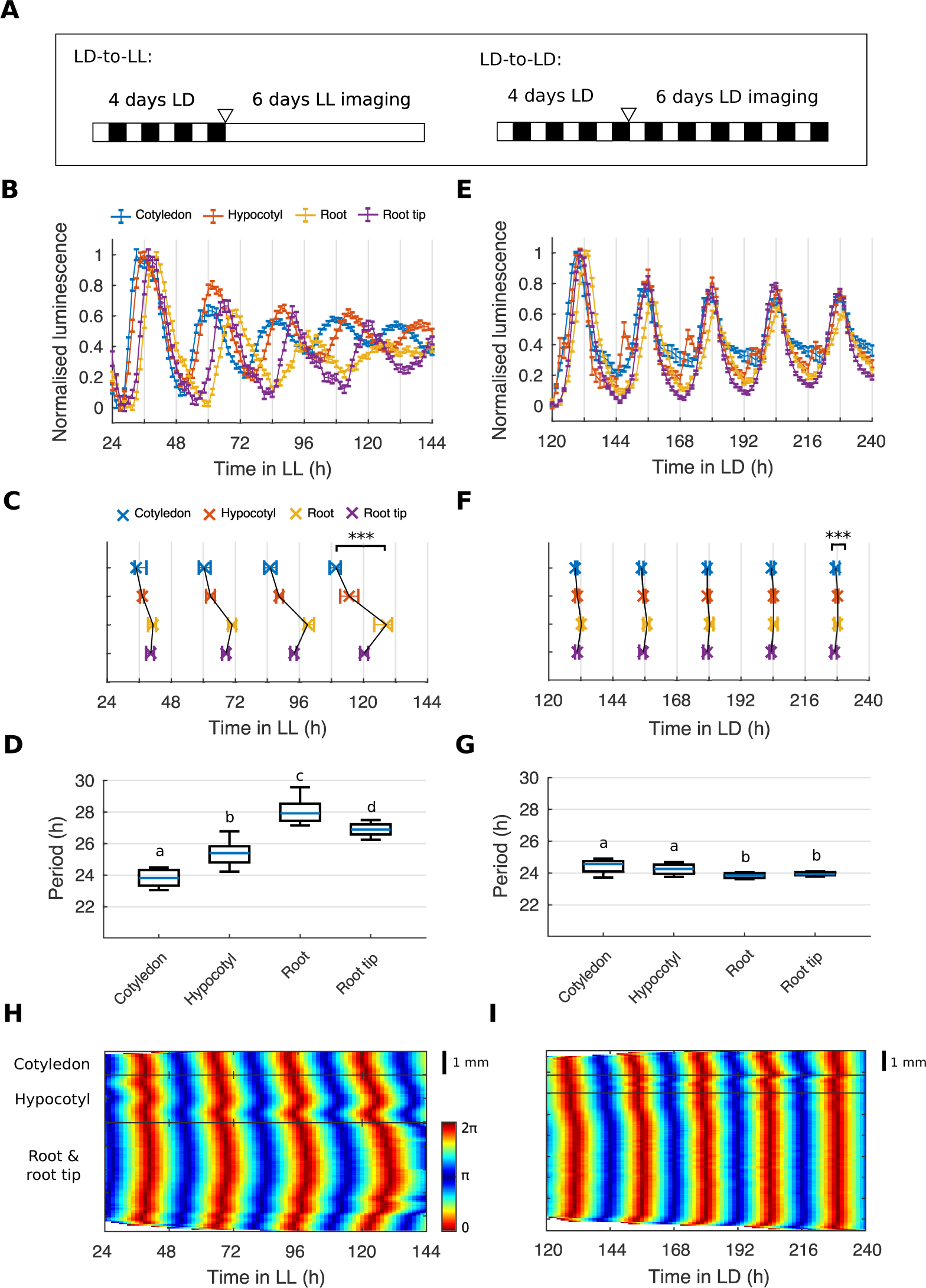
Organ specific clocks show phase differences under constant environmental conditions and light-dark cycles. A. Schematic depicting the experimental conditions used. Seedlings were grown for 4 days under light-dark (LD) cycles and imaged either under constant light (LD-to-LL) or LD (LD-to-LD). The white triangle represents the beginning of imaging. B. Expression of *GI::LUC* from different organs imaged under the LD-to-LL condition. Luminescence counts were normalised to the minimum and maximum value of the time-series. Data represents the mean ± S.E.M. of all rhythmic time-series. C. Times of peaks of expression in different organs under LD-to-LL condition. Plots represent the 25^th^ percentile, median, and the 75^th^ percentile for the peak times of the oscillations of each tissue. Organs show significant phase differences, *** *p* < 0.001, by Kruskal-Wallis ANOVA. Pairwise comparisons are shown in S1 Fig. D. Period estimates for different organs imaged under LD-to-LL condition. The means of organs are statistically different (*p* < 0.05, by one-way ANOVA, Tukey’s *post ho*c tests) if they do not have a letter in common. E. Expression of *GI::LUC* from different organs imaged under the LD-to-LD condition. Luminescence counts were normalised to the minimum and maximum value of the time-series. Data represents the mean ± S.E.M. of all rhythmic time-series. Color legend is as in B. F. Times of peaks of expression in different organs imaged under LD-to-LD condition. Plots represent the 25^th^ percentile, median, and the 75^th^ percentile for the peak times of the oscillations of each tissue. Organs show significant phase differences, *** *p* < 0.001, by Kruskal-Wallis ANOVA. Pairwise comparisons are shown in S1 Fig. Color legend is as in C. G. Period estimates for different organs imaged under LD-to-LD condition. The means of organs are statistically different (*p* < 0.05, by one-way ANOVA, Tukey’s *post ho*c tests) if they do not have a letter in common. H, I. Representative phase plot of *GI::LUC* expression across longitudinal sections of the cotyledon (top), hypocotyl (middle), and root (bottom) of a single seedling under LD-to-LL (A) and LD-to-LD (B) condition. Colorbars are as in H. For LD-to-LL data, *N* = 4; LD-to-LD, *N* = 3; For both, *n* 25. *N* represents the number of independent experiments, *n* the total number of seedlings. See S1 and S2 File for exact *n* and test statistics. All boxplots indicate the median, upper and lower quartile, and whiskers the 9^th^ and 91^st^ percentile.

The phase at which a rhythm entrains to the environment can depend on the mismatch between its endogenous period and the period of the entraining signal (27–29). We therefore tested the consequence of endogenous period differences between organs on the entrainment of the plant, by monitoring rhythms under LD cycles (LD-to-LD; Fig 2A and Materials and Methods). Under the LD-to-LD condition, we observed robust and entrained rhythms of *GI*::*LUC* (Fig 2E). However, closer inspection of the timing of the peaks of the oscillations revealed significant differences in clock phase between organs (Fig 2F). The cotyledon and hypocotyl consistently peak earlier than the root regions, but the root tip peaks earlier than the middle of the root (Fig 2F, S1 Fig and S2 Video). This is qualitatively similar to the pattern observed under LL (Fig 2C). However, under the LD-to-LD condition, the organs showed a more stable phase relationship than under LL, with a range between the earliest and latest peaking region of 2.08 ± h in the first and 1.10 ± 1.44 h in the final oscillation. This is due to the fact that all organs oscillate with a period of approximately 24 h (Fig 2G).

### Spatial waves of clock gene expression propagate between and within organs

Spatial waves of clock gene expression have been previously reported in plant leaves (14,15,22,23) and roots (5,16,24) under LL. However, their relation to one another, and the relevance under LD cycles remained unclear. We analyzed our LD-to-LL and LD-to-LD dataset of whole, intact seedlings at the sub-tissue level in order to address these questions. We extracted the phase of the luminescence signal across longitudinal sections of seedlings (S2 Fig, Materials and Methods) and present phase plots and time-lapse videos of single seedlings representative for each light condition (Fig 2H, I, and S1,2 Video). The clearest waves of expression could be observed in the LD-to-LL condition, as phase differences increased with time. In the cotyledon, a wave of *GI*::*LUC* expression propagated from the tip to the base (Fig 2H, top), and downwards into the hypocotyl (Fig 2H, middle). In the hypocotyl we observed a second wave traveling from the root junction upwards into the hypocotyl (Fig 2H, middle). Finally, within the root we observed two waves; one propagating down from the hypocotyl junction and the second from the root tip upwards into the root, as we have reported previously (Fig 2H, bottom; 5). Evidence of waves of clock gene expression could also be observed under the LD-to-LD condition. Although they are less pronounced, small phase waves could be discerned within the cotyledon (Fig 2I, top), hypocotyl (Fig 2I, middle) and root (Fig 2I, bottom) of the phase plots and time-lapse videos (S2 Video).

### Spatial waves of clock gene expression persist in the absence of inter-organ communication

Previous work has proposed that spatial waves of clock gene expression are driven by local cell-cell coupling (5,14–16). However, plants can communicate through both local and long-distance, inter-organ pathways (30), and the root clock has been proposed to be driven by long range signals from the shoot (18). To investigate whether rhythms and spatial waves are driven by long-distance communication we blocked signal transmission between organs by cutting the seedling into sections. We cut the root at either the hypocotyl junction, the root tip, or both the hypocotyl junction and the root tip, and then monitored the rhythms for six days under LL (Fig 3A). Surprisingly, we found that sectioning the plant did not significantly affect the phase of the rhythms (Fig 3B–D and S3 Fig). We found that this is due to the persistence of period differences across the plant after cutting (Fig 3E–G). Next, we focused our analysis to within the hypocotyl and root, where the simple geometry means the wave patterns can be most easily observed. Strikingly, after all cuts we observed the persistence of waves propagating both from the hypocotyl down into the root and from the root tip upwards (Fig 3I–K and S3 Video). Our results show that in all organs excised, rhythms are autonomous and the spatial waves that travel between them are not dependent on a long-distance signal.

**Fig 3.**
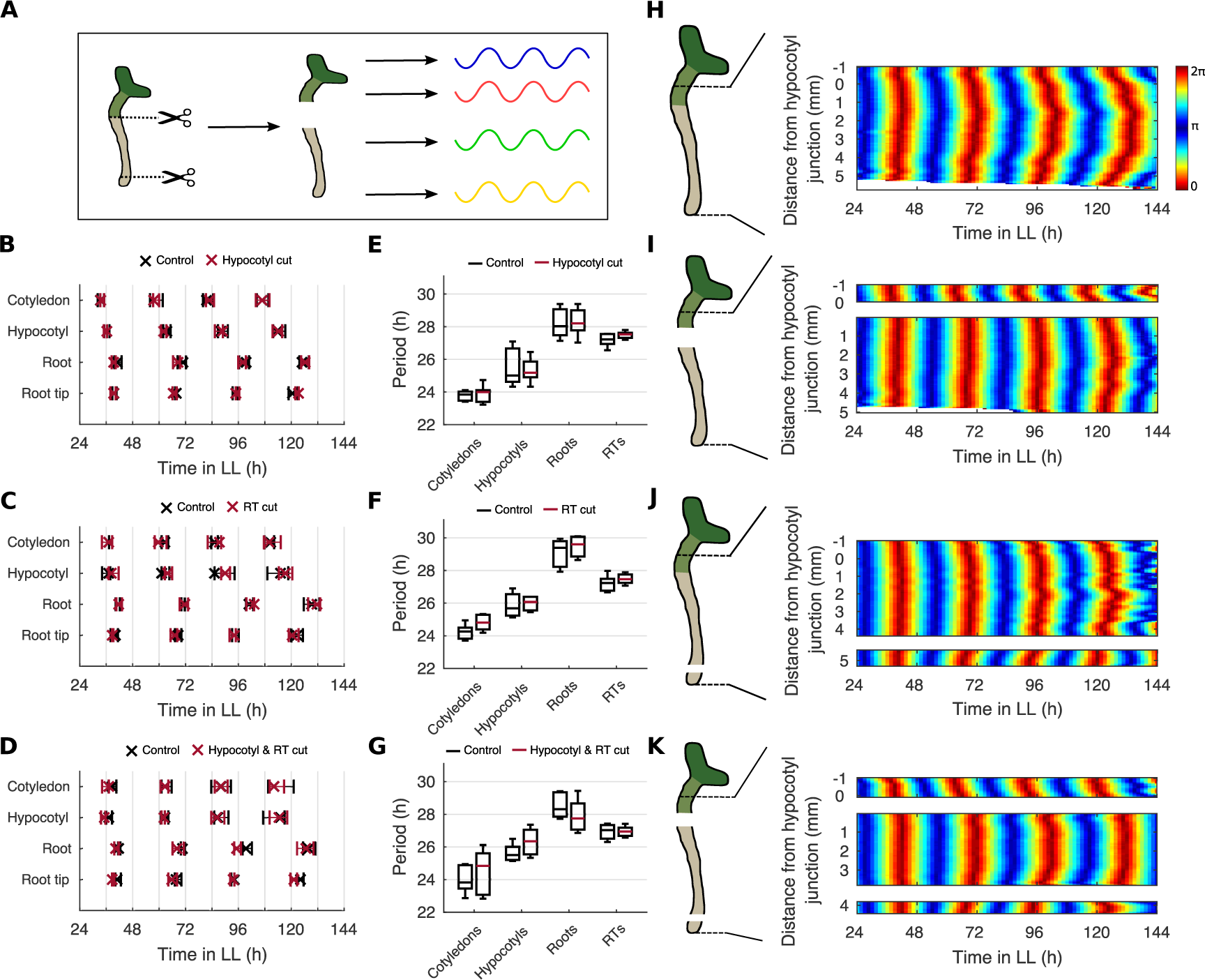
Spatial waves of clock gene expression persist in the absence of long-distance signals. A. Schematic depicting the experimental design. Seedlings were cut at the hypocotyl junction, root tip (RT), or at both the hypocotyl junction and the RT. The rhythm of both the excised organs and the remaining intact organs were subsequently analyzed. B-D. Times of peaks of expression in different organs following a cut at the hypocotyl junction (B), RT (C), or both the hypocotyl junction and RT (D). Plots represent the 25^th^ percentile, median, and the 75^th^ percentile for the peak times of the oscillations of each tissue. E-G. Period estimates for different organs following a cut at the hypocotyl junction (E), RT (F), and both the hypocotyl junction and RT (G). All comparisons of means are not significantly different, *p* > 0.05, by two-tailed *t*-test, Welch correction. H-K. Representative phase plot of *GI::LUC* expression across longitudinal sections of the hypocotyl and root of a single seedling without a cut (H) or with a cut at either the hypocotyl junction (I), RT (J), or both the hypocotyl junction and RT (K). Schematic shows the approximate cut position and the region analyzed. Colormaps are as in H. For hypocotyl cut experiments, *N* = 4; root tip cut, *N* = 4; hypocotyl and root tip cut, *N* = 4. For all, *n* 13. *N* represents the number of independent experiments, *n* the total number of seedlings. See S1 and S2 File for exact *n* and test statistics. All boxplots indicate the median, upper and lower quartile, whiskers the 9^th^ and 91^st^ percentile.

### Period differences plus local coupling can explain organ specific entrainment and spatial waves

The persistence of rhythms and spatial waves in the absence of long-distance communication suggests clocks may instead be coupled through local interactions. We extended the mathematical framework we employed in Gould *et al*, 2018 to investigate whether local coupling can explain the entrainment behaviors that we observe under LD and LL. As before, we used a Kuramoto phase oscillator model (27). In this framework each pixel (which in fact represents multiple cells (S4 Fig)) on our seedling template is an individual oscillator with an intrinsic period and is weakly coupled to its nearest neighbors. The intrinsic period of each pixel is set according to its location in the seedling. Pixels from the cotyledon, hypocotyl, root, and root tip were drawn from distributions centered around the mean periods that we observed experimentally in each region under LL (Fig 4A, S4 Fig, Materials and Methods). These period estimates are made from *in vivo* experiments and therefore include the effects of coupling. They are, however, as good an estimation of the cell autonomous periods as possible in a physiologically relevant context. In our LD-to-LL simulations, due to the differences in intrinsic periods, and coupling, we see increasing phase shifts between organs (Fig 4C), and two increasingly large waves in the root (Fig 4E), as observed in experiments (Fig 4G).

**Fig 4.**
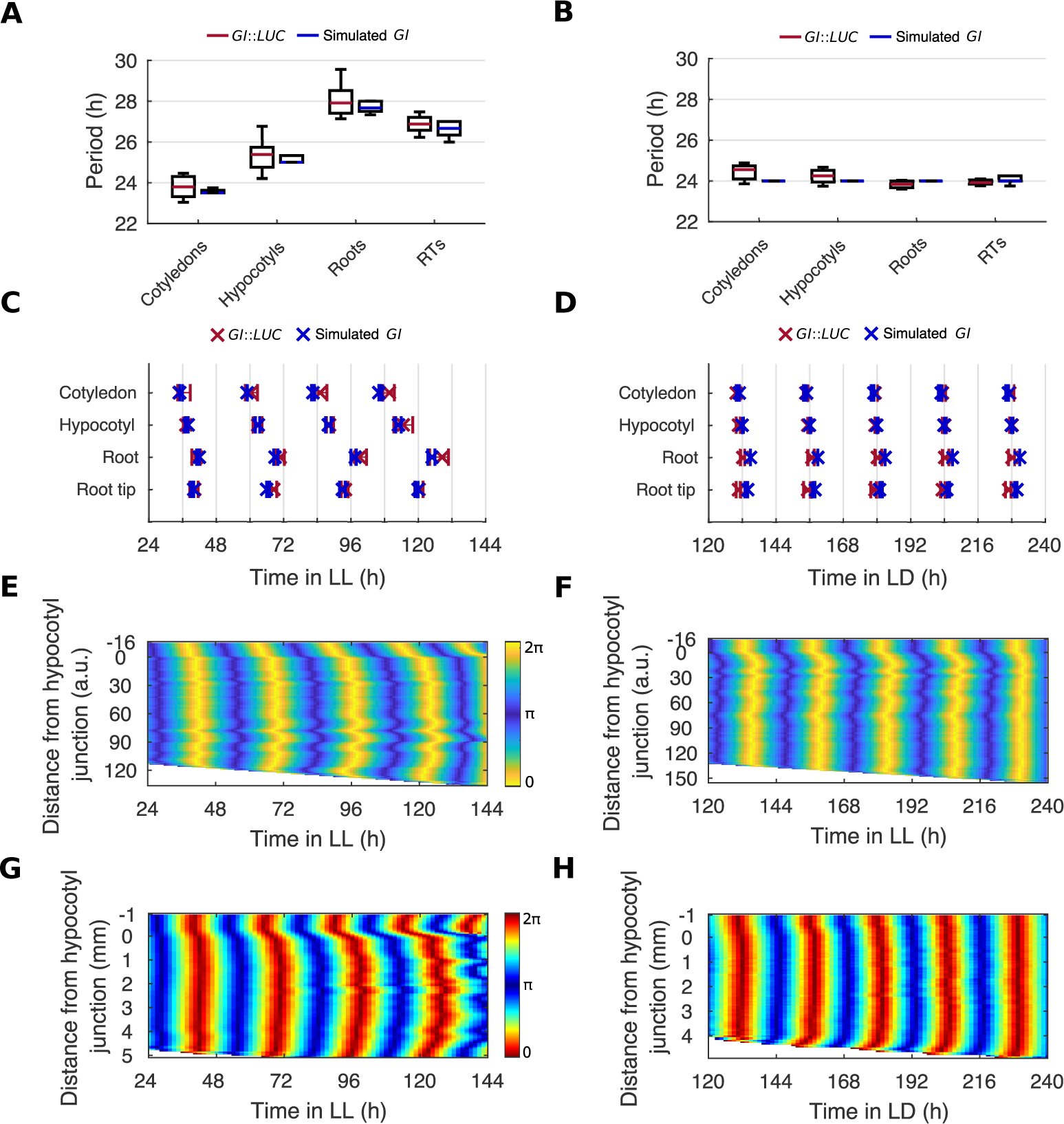
Period differences and local coupling can explain spatial waves of clock gene expression. A, B. Period estimates of simulated *GI* for different organs imaged under LD-to-LL (A) and LD-to-LD (B) condition. C, D. Times of peaks of expression for simulations and experimental data in different organs under LD-to-LL (C) or LD-to-LD (D) conditions. Plots represent the 25^th^ percentile, median, and the 75^th^ percentile for the peak times of the oscillations of each tissue. E, F. Representative phase plot of simulated *GI* expression across longitudinal sections of the hypocotyl and root of a single seedling under LD-to-LL (E), or LD-to-LD (F) conditions. Colormaps are as in E. G, H. Representative phase plot of *GI::LUC* expression across longitudinal sections of the hypocotyl and root of a single seedling under LD-to-LL (G) and LD-to-LD (H) conditions. Colormaps are as in G. For experimental data *N* and *n* are as in Fig 2. For simulations, *n* = 24. *N* represents the number of independent experiments, *n* the total number of seedlings. See S1 and S2 File for exact *n* and test statistics. All boxplots indicate the median, upper and lower quartile, whiskers the 9^th^ and 91^st^ percentile.

In our model, the amount that each oscillator phase is shifted is set by the mismatch of its intrinsic period and the period of the entraining rhythm (27–29). This prediction is supported by experimental evidence in various organisms, including plants (31), although dawn can also reset the phase of the plant clock in bulk *Arabidopsis* experiments (32). We tested whether the phase differences that we observe between organs in *Arabidopsis* under our LD conditions can be reproduced in our model by this mismatch with the entraining rhythm. In our simulations, organs were forced to oscillate with a period of approximately 24 h, due to entrainment to the external rhythm (Fig 4B). However, due to the mismatch between the intrinsic period and the entraining rhythm, organs entrained with different phases, matching those observed experimentally (Fig 4D). Phase shifts could also be observed at the sub-tissue level; two short waves could be observed in the root (Fig 4F), as in experiments (Fig 4H).

### Local coupling limits desynchrony in the absence of entrainment

In a set of coupled oscillators, variation in period causes a decrease in synchrony, whereas coupling and external entrainment maintain or increase synchrony (33,34). In order to make predictions about the presence of local coupling in seedlings we simulated our model in the absence of LD entrainment. We simulated the duration of the experiment without entraining the oscillators, and thus assume that the phases are initially random (LL-to-LL; Fig 5A, Materials and Methods). In contrast to the LD-to-LL condition, where oscillators begin synchronous but become less synchronized whilst under LL, in LL-to-LL simulations, oscillators began less synchronous but maintained their order over the six days (Fig 5B). Interestingly, in the root, the model predicted a complex spatial pattern, with multiple phase clusters and spatial waves in a single seedling (Fig 5C and S4 Video). These patterns of gene expression were similar to the zig-zag patterns previously reported by others when roots are grown on sucrose supplemented media (16,24,35). We found that these zig-zag patterns emerged with, but not without, local coupling (S5A and S5B Fig). Simulations of a plausible alternative model without coupling but with a gradient of the intrinsic periods was sufficient to generate the simple waves that we observed under LD-to-LL (S5C and S5D Fig), but not the complex zig-zag waves predicted in the LL-to-LL condition (S5E and S5F Fig).

**Fig 5.**
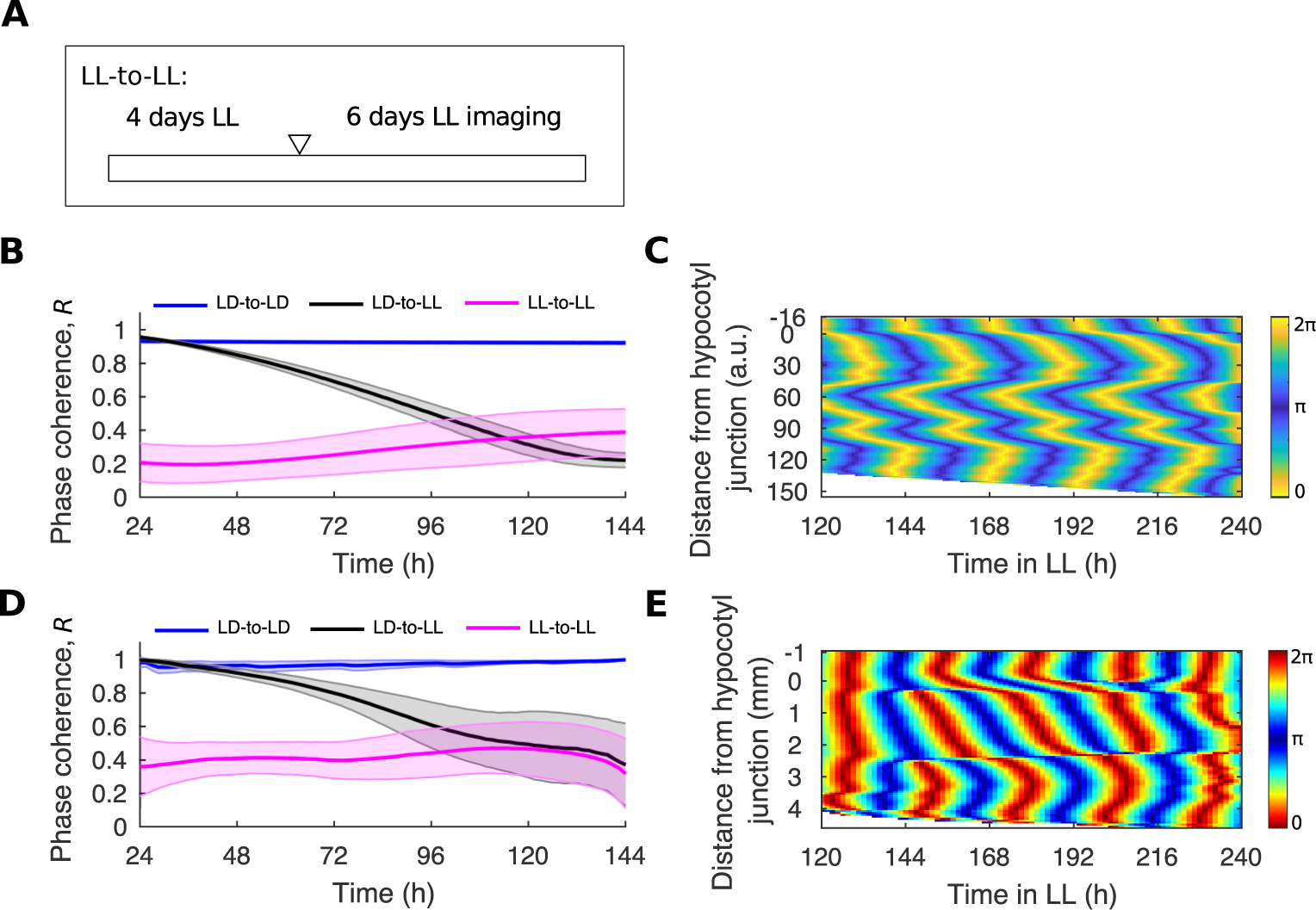
Local coupling limits desynchrony in the absence of light-dark cycles. A. Schematic depicting the experimental conditions used. Seedlings were grown for 4 days under LL and then imaged also under LL (LL-to-LL), so that seedlings have never seen an LD cycle. The white triangle represents the beginning of imaging. B. Quantification of phase coherence by time evolution of the Kuramoto order parameter, *R*, for simulated *GI* expression. Solid lines indicate the mean and the shaded region one S.D. of the mean. C. Representative phase plot of simulated *GI* expression across longitudinal sections of the hypocotyl and root of a single seedling under LL-to-LL condition. D. Quantification of phase coherence by time evolution of the Kuramoto order parameter, *R*, for *GI::LUC* expression. Solid lines indicate the mean and the shaded region one S.D. of the mean. E. Representative phase plot of *GI::LUC* expression across longitudinal sections of the hypocotyl and root of a single seedling under the LL-to-LL condition. For *GI* model simulations, *n* = 24; for LL-to-LL *GI::LUC* data, *N* = 3 and *n* 25. *N* represents the number of independent experiments, *n* the total number of seedlings. See S1 and S2 File for exact *n* and test statistics.

In order to test our model and validate the assumption of local coupling, we experimentally tested the LL-to-LL model prediction. We both grew and imaged seedlings under LL conditions (LL-to-LL; Fig 5A), so that seedlings never see an entrainment cue beyond germination (36,37). Roots maintain their coherence over the six days of imaging (Fig 5D) and display a zig-zag expression pattern (Fig 5E and S6 Fig) as predicted by the model, supporting the hypothesis of weak, local coupling.

### Local light inputs set organ specific periods

To test our model further, we attempted to manipulate the periods in specific organs, to determine whether we could modulate the spatial waves of gene expression. In the most severe case, removing all period differences across the plant should result in perfectly coherent rhythms. We found mutations to the core clock network to have little effect on the organ specificity of periods (S7 Fig), and so we next tested whether we could alter periods in an organ specific manner by modulating inputs to the clock. We first tested the effect of light input, by growing seedlings under LD cycles before imaging seedlings under constant darkness (DD). Under DD we observed a drastic slowing of periods in the cotyledon and hypocotyl but an increase in speed at the root tip (Fig 6A). This caused a reduction of phase shifts between the aerial organs and the root (Fig 6B and S8 Fig), and the loss of spatial waves traveling from the hypocotyl down the root (Fig 6C and S5 Video). Inversely, the faster periods at the root tip caused a larger phase shift between the root tip and the root (Fig 6B and S8 Fig), resulting in a longer spatial wave traveling from the root tip upwards into the root (Fig 6C and S5 Video). We observed the same effect when seedlings were grown hydroponically, so that roots did not see light during entrainment or imaging (S9A–C Fig). Additionally, a qualitatively similar but lesser effect was observed under monochromatic red or blue light (S9D–F Fig).

**Fig 6.**
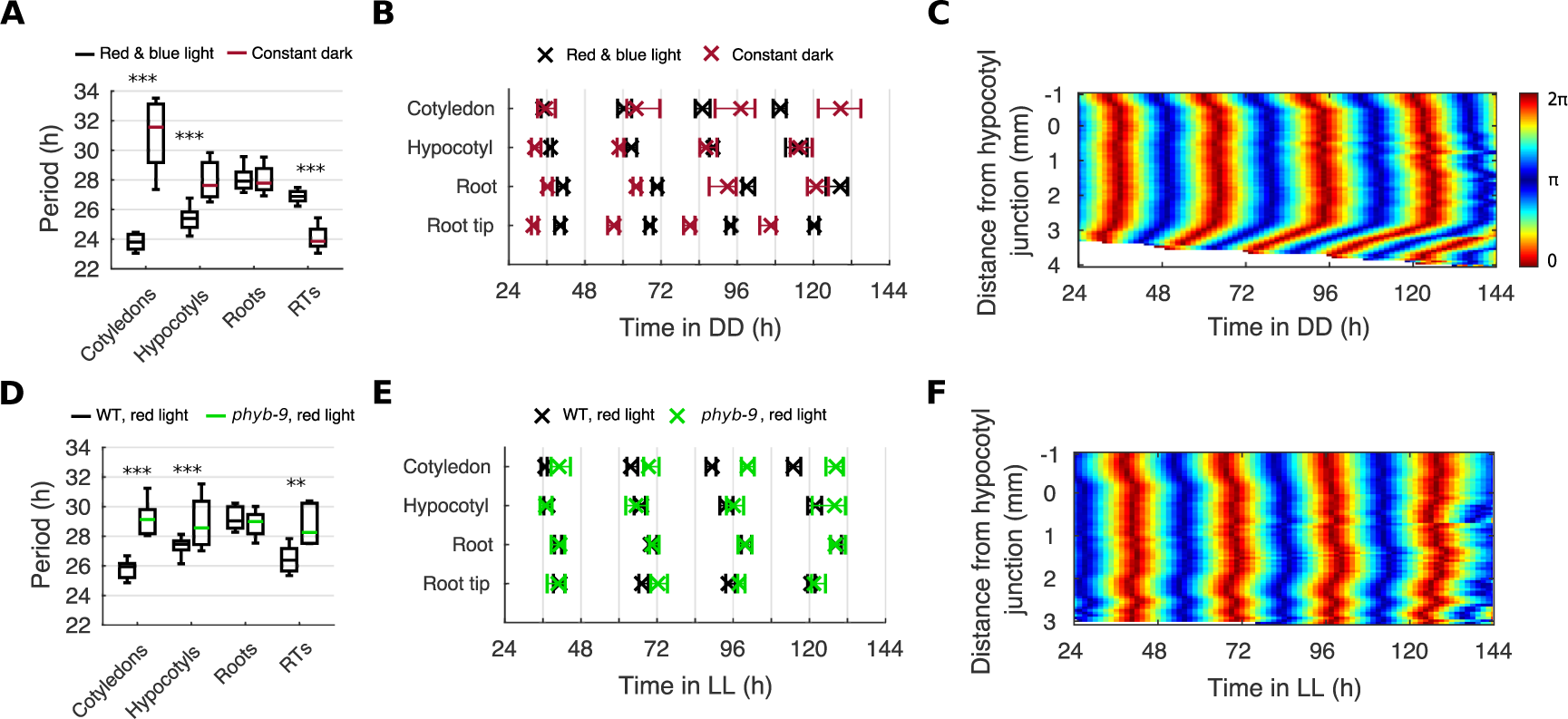
Light input sets the period of clocks organ specifically. A. Period estimates for different organs under constant red and blue light or constant darkness (DD). *** *p* < 0.001, by two-tailed *t*-test, Welch correction. B. Times of peaks of expression in different organs under constant red and blue light or constant darkness. Plots represent the 25^th^ percentile, median, and the 75^th^ percentile for the peak times of the oscillations of each tissue. C. Representative phase plot of *GI::LUC* expression across longitudinal sections of the hypocotyl and root of a single seedling under DD. D. Period estimates for different organs under constant red light in the *phyb-9* mutant. ** *p* < 0.01, *** *p* < 0.001, by two-tailed *t*-test, Welch correction. E. Times of peaks of expression in different organs under constant red light in the *phyb-9* mutant. Plots represent the 25^th^ percentile, median, and the 75^th^ percentile for the peak times of the oscillations of each tissue. F. Representative phase plot of *GI::LUC* expression across longitudinal sections of the hypocotyl and root of a single seedling under constant red light in the *phyb-9* mutant. For constant red & blue light, *N* = 4; DD, *N* = 3; *phyb-9, N* = 4. For all, *n* 25. *N* represents the number of independent experiments, *n* the total number of seedlings. See S1 and S2 File for exact *n* and test statistics. All boxplots indicate the median, upper and lower quartile, and whiskers the 9^th^ and 91^st^ percentile. Red & blue light data is a re-plot of LD-to-LL data from Fig 2, for comparison.

We next tested whether the effect of light on organ specificity is direct, through known light signaling pathways. We imaged *GI*::*LUC* expression in the *phyb-9* background, a null mutant for the primary red light photoreceptor in *A. thaliana*, PHYTOCHROME B (38,39). Under red light, in the *phyb-9* mutant we observed the loss of period differences between the cotyledon, hypocotyl, and root (Fig 6D). This caused the loss of phase shifts between the aerial organs and the root (Fig 6E), and the loss of spatial waves traveling down the root (Fig 6F and S6 Video). We also observed a decrease in rhythmicity across the seedling (S1 File). The effect was particularly large in the root tip, with only 24 % of root tips classed as rhythmic compared to 96 % in the wild type. In the root tips classed as rhythmic, the period ran approximately 3 h slower, at approximately the same speed as the middle of the root (Fig 6D). Therefore after six days, in all seedlings, the phase shift between the root tip and root (Fig 6E and S10A Fig), and the spatial wave traveling from the root tip upwards, was lost (Fig 6F and S6 Video). The *phyb-9* mutation, however, does not abolish the faster periods observed in the root tip under constant darkness (S10B, C Fig).

### Local metabolic inputs set organ specific periods

In addition to the external environment, the circadian clock is exposed to biochemical signals from within the cell (40). We investigated whether these endogenous signals could also alter periods in an organ specific manner, modulating the spatial waves of clock gene expression. First, we imaged seedlings under LL in the presence of 3-(3,4-dichlorophenyl)-1,1-dimethylurea (DCMU), a specific inhibitor of photosynthesis. During inhibition, we observed a slowing of periods specifically in the cotyledon and hypocotyl (Fig 7A), causing a loss of phase shifts between the hypocotyl and root (Fig 7B and S11 Fig), and the loss of spatial waves down the root (Fig 7C).

**Fig 7.**
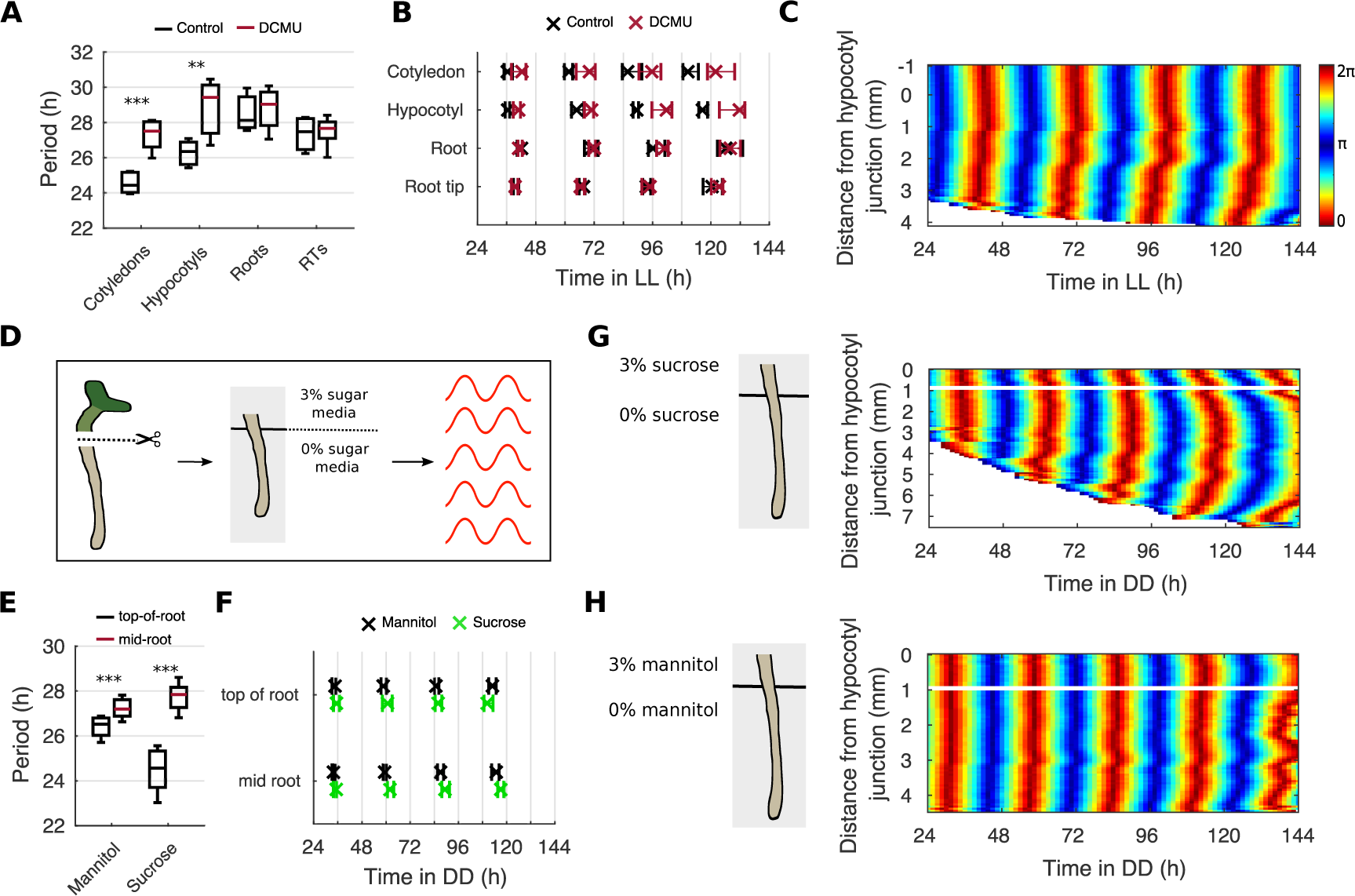
Photosynthetic sugar sets the period of clocks organ specifically. A. Period estimates for different organs during the inhibition of photosynthesis by DCMU. ** *p* < 0.01, *** *p* < 0.001, by two-tailed *t*-test, Welch correction. B. Times of peaks of expression in different organs during the inhibition of photosynthesis by DCMU. Plots represent the 25^th^ percentile, median, and the 75^th^ percentile for the peak times of the oscillations of each tissue. *** *p* < 0.001, by Kruskal-Wallis ANOVA. Color legend is as in A. C. Representative phase plot of *GI::LUC* expression across longitudinal sections of the hypocotyl and root of a single seedling during the inhibition of photosynthesis by DCMU. D. Schematic representing the experimental design. Seedlings are cut at the hypocotyl junction and the excised root laid across two adjacent agar pads, one containing sugar supplemented media and the other not, so that only the top part of the root is in contact with sugar. Roots are then imaged under constant darkness. E. Period estimates for the top and middle region of the root during the partial contact of the root with sucrose or mannitol, under constant darkness. *** *p* < 0.001, by two-tailed *t*-test, Welch correction. F. Times of peaks of expression for the top and middle region of the root during the partial contact of the root with exogenous sucrose or mannitol, under constant darkness. G, H. Representative phase plot of *GI::LUC* expression across longitudinal sections of the hypocotyl and root of a single seedling during the partial contact of the root with exogenous sucrose (G) or mannitol (H), under constant darkness. Schematic shows the approximate root positioning on the agar pads. Colorbars are as in C. For DCMU, *N* = 3; exogenous sugar experiments, *N* = 3; For all, *n* 25. *N* represents the number of independent experiments, *n* the total number of seedlings. See S1 and S2 File for exact *n* and test statistics. All boxplots indicate the median, upper and lower quartile, and whiskers the 9^th^ and 91^st^ percentile.

Photosynthesis modulates the clock through the production of sugars, which feed into the oscillator (41–43). We next tested whether the application of sucrose to part of the plant could locally reduce clock periods and generate a spatial wave. This is a direct test of the hypothesis that local period differences drive spatial waves of gene expression. We designed a protocol that allowed us to rest only the top portion of the root on sugar supplemented media, and observe the effect throughout the root. We did this with roots cut at the hypocotyl junction, to minimize developmental effects, and under DD, where we ordinarily observe no spatial wave down the root (Fig 7D). In comparison to mannitol (a poorly metabolized sugar that acts as an osmotic control), contact with sucrose supplemented media caused a larger decrease in period length (Fig 7E). This caused a larger phase shift from the top to the middle of the root (Fig 7F and S12 Fig). Within the root, a clear spatial wave of clock gene expression propagates down from the top of the root when in contact with the sucrose (Fig 7G and S7 Video), but not mannitol (Fig 7H and S8 Video) supplemented media. Together these results show that speeding up clocks locally, either via modulating light perception or the addition of photosynthetic sugars, can drive spatial waves of clock gene expression.

## Discussion

Here, we report how local periods, due to differences in sensitivity to clock inputs, generate spatial waves of circadian clock gene expression across the plant. Using time-lapse imaging we show that spatial waves exist both in constant and entrained conditions and do not require long distance signals. Modeling and experiments show that local coupling can explain our results, including complex synchronization patterns in plants that have never seen an entraining signal. Finally, by the manipulation of environmental inputs, we are able to modulate the waves in a predictable manner by locally altering clock periods. We therefore propose that spatial waves are sufficient to integrate organ specific environmental inputs and coordinate timing across the plant.

In the laboratory, clocks are most often studied under constant environmental conditions in order to observe the endogenous genetic properties of the oscillator. However, in the wild, plants are exposed to environmental cycles and the interaction between the oscillator and the environment is of importance. It is therefore significant that we observed phase differences between clocks within a plant, even under LD cycles. A previous high resolution study in *A. thaliana* observed phase differences within leaves after the transfer from LL to LD conditions, though rhythms were near synchronous after three days in LD cycles (14). Phase differences have also been observed in *lemna gibba* fronds, where cells in leaves entrain with different phases, causing a centrifugal pattern (17). Phase patterns under LD cycles therefore appears to be a common property of plant circadian systems, and will require further investigation.

The presence of local cell-cell coupling has been previously suggested to help maintain clock synchrony within *A. thaliana* (5,14–18). In addition, long-distance signals (18,21), and light piped from the shoot (25), have been proposed as mechanisms for coordination. Through a combination of experiments and modeling we show that in seedlings, local signals alone are sufficient to maintain robust rhythms over six days in all organs, as well as generate the observed complex spatial patterns in clock expression. We note that our results do not exclude the possibility that phloem mobile signals, or light piped from the stem, additionally act to synchronize the root with the shoot. However, the waves that we observe in cut roots, combined with the wave up the root apparent in seedlings grown in constant darkness, suggests that these signals do not drive the spatial wave patterns that we observe. In future work it will be important to investigate whether coordination through local coupling also occurs in later stages of plant development, and if so, whether the coordination structure changes as the plant develops to compensate for its increasing size.

Local coupling is dependent on a signal that is cell-to-cell mobile. Research in cellular communication in plants has intensified in recent years and a number of signals are known to be mobile between cells and tissues. A selection of hormones, sugars, mRNA’s, proteins, and ions have been shown to be both mobile, and capable of influencing the clock (3,40). To better understand the mechanism of intercellular coupling of clocks in plants it will be important to investigate whether one, some, or all of these mobile signals act to couple the clock. The study of this will benefit greatly from the development of ‘omics’ methods at the single cell level (44,45).

Oscillators in different organs of the plant will be exposed to different environments, both externally from the environment and internally from the cell’s biochemistry. We found that these differences in input can drive spatial waves by creating period differences. We demonstrated this by manipulating two environmental inputs, light and sucrose, an external and internal signal respectively. Light intensity is transmitted to the clock by phytochromes and cryptochromes, causing a decrease in period (46,47). Since these genes have tissue specific expression patterns in the plant (48–51), we can modulate the periods locally using light. We did this by controlling the quality of the light or by perturbing light signaling using a *phyb-9* genetic background. In both cases we successfully modulated spatial waves of clock gene expression, and were able to abolish them in the *phyb-9* background under red light, due to the minimization of period differences across the plant. In a similar fashion, by perturbing photosynthesis or by directly applying exogenous sucrose to roots, we found that we can affect periods locally and modulate spatial waves of clock gene expression. There are, however, many other signals known to modulate the speed of the clock (40). In future work it will be important to test how these interact, and the consequence to spatial coordination when plants are under physiological conditions. Of particular interest will be temperature, which is known to differ between the air and the ground (52) and deviate from the photoperiod (53). In fact, it has already been demonstrated that temperature is preferentially sensed by the clock in specific cell types (10,11). Comprehensive *in vivo* studies, under a range of environmental conditions, will be required to understand the full complexity.

For plants, being responsive to the environment whilst being robust to fluctuations necessitates a trade-off. The clock, in its role as master regulator, must balance these two competing requirements. Recently it has been proposed that the clock in plants is dynamically plastic, able to respond to changes in environmental inputs by altering phase and period (54,55). A decentralized structure, with organ specific inputs to clocks that are coupled together, could allow some flexibility in sensing the environment whilst ensuring robust timing. In future, it will be important to better understand the importance of this design principle to physiological outputs of the clock and the development of the plant.

## Materials and Methods

### Plant materials and growth conditions

The wild type *GI*::*LUC* line is in the Col-0 background and as described previously (56). The *cca1-11* (TAIR:1008081946; 57; Ws background back-crossed with Col-0 three times), *prr9-1* (TAIR:3481623; 55), *prr7-3* (TAIR:3662906; 55), *toc1-101* (TAIR:6533848449; 56), and *lux-4* (TAIR:1008810333; 58) alleles are loss of function mutations that have been previously described, and were transformed with the *GI*::*LUC* (56) construct by means of *Agrobacterium* mediated transfection (61).

Seeds were surface sterilized and placed in the dark at 4 °C for 3 days. Seeds were sown at dawn of the fourth day on full strength Murashige & Skoog (MS), 2 % agar, pH 5.7 media, without sucrose unless otherwise specified. Seeds were then grown inside of plant growth incubators (MLR-352; Panasonic, Japan) for 4 days under 80 mmol m^2^ s^−1^ cool white light at a constant temperature of 22 °C. Seedlings were grown under 12 h light-12 h dark cycles unless otherwise specified. Plates were orientated vertically during growth.

For experiments where roots are grown in the dark (S9 Fig), seedlings were grown hydroponically in full strength MS liquid solution as described previously (62). After four days of growth, working under green light only, seedlings were transferred to MS 2 % agar plates and transferred to imaging cabinets.

### Luciferase imaging

At dusk of the fourth day of growth, seedlings were sprayed with a 5 mM D-Luciferin (Promega, USA), 0.01 % Triton X-100 solution. At dawn of the fifth day, 6–8 seedlings were transferred into a 3-by-3 cm area of a media plate in order to fit inside of the camera’s field of view. Plates were orientated vertically during imaging.

Imaging was performed inside of growth incubators (MIR-154; Panasonic, Japan) at a constant temperature of 22 °C and under an equal mix of red and blue light emitting diodes (40 μmol m^−2^ sec^−1^ total), unless specified as red light only (RLL; 40 μmol m^−2^ sec^−1^ red) or blue light only (BLL; 40 μmol m^−2^ sec^−1^ blue). For experiments under LD cycles, lights were switched on to full intensity at dawn and completely off at dusk. Images were taken every 90 minutes for six days, with an exposure time of 20 minutes. Images were taken using a LUMO (QImaging, Canada) charge-coupled device (CCD) camera, controlled using micro-manager (V2.0; Open Imaging) as previously described (63). The camera lens (Xenon 25 mm f/0.95; Schneider, Germany) was modified with a 5 mm optical spacer (Cosmicar, Japan) to increase the focal length and decrease the working distance.

### Cuts and treatments

For cut experiments seedlings were cut approximately 3 h after dawn of the fifth day of growth, immediately prior to the commencement of imaging. For ‘hypocotyl cut’ experiments (Fig 3B, E, and I) seedlings were cut in the root as close to the hypocotyl junction as discernible by eye, for ‘root tip cut’ experiments (Fig 3C, F and J) seedlings were cut approximately 100–200 μm from the root cap. Cuts were made with a pair of Vanna’s type microdissection scissors (Agar Scientific, UK). Following all excisions, the organs were gently separated with a pair of forceps to ensure no physical contact.

DCMU was added to the media at a final concentration of 20 mM. Seedlings were transferred to the DCMU containing media at dusk of the fourth day of growth. For sugar application experiments (Fig 7E–H), media was added in 8-well rectangular dishes (NUNC; Thermo-Fisher Scientific) so that one well contains media supplemented with MS and sugar whilst the adjoining well contains media supplemented with MS only. Wells were filled with equal volumes to the brim of the wells so that the two agar pads form a continual flat surface but do not touch. Sucrose or mannitol was added at a final concentration of 90 mM (3 % w/v). Seedlings were cut at the hypocotyl junction (as described above), and laid across the adjoining agar pads so that approximately the top 1 mm of the excised root rests on the sugar supplemented media, and the remainder of the root rests on the non-sugar supplemented media. Seedlings were cut and transferred to the media at dawn of the fifth day of growth, immediately prior to the commencement of imaging.

### Organ level analysis of period and phase

For the organ level analysis of the period and phase, organs were first tracked manually in Imaris (BitPlane, Switzerland) using the ‘Spots’ functionality. We use a circular region of interest (ROI) of approximately 315 mm diameter and track the center of a single cotyledon, hypocotyl, root, and the root tip from each seedling. As the root grows we maintain the root ROI a fixed distance from the hypocotyl junction. A small number of cotyledons and hypocotyls were not trackable due to their orientation or their overlap with each other. These organs were excluded from the analysis. The median of the ROI was extracted to give the time-series. Prior to the analysis of period and phase, the time-series were first background subtracted. Very low expression rhythms with a minimum intensity value of less than zero after background subtraction were then removed. All time-series were inspected by eye after pre-processing steps and prior to analysis.

Period analysis was conducted in BioDare2, a data server for the analysis of circadian data (biodare2.ed.ac.uk; 61). All period estimates were performed on non-normalized data between 24–144 h from dawn of the day imaging began using the Fast-Fourier Transformed Non-Linear Least Squares (FFT-NLLS) algorithm (65,66). Data was first baseline detrended by subtraction of a polynomial of degree three from the data. Oscillations were classed as rhythmic if the FFT-NLLS algorithm returned a period in the range of 18–36 h with a confidence level (as defined in (61)) below 0.6.

For the analysis of the times of peaks of expression, peaks were identified using the MATLAB ‘findpeaks’ function. This was done after the application of a third order Butterworth filter to remove high frequency noise. Only peaks where all organs complete the full cycle within 24–144 h from dawn of the day imaging are used. Additionally, peaks were discarded if they are closer than 18 h or further than 36 h apart.

### Statistical analyses

In all figures data points, measure of error, statistical test used, *n*, and approximate *p* values are reported in the figure legend. Exact *p* values, exact *n*, and other test statistics are reported in S1 and S2 File. When values are described in the text, they are quoted as mean ± standard deviation of the mean. For the comparisons of period estimates, one-way analysis of variance (ANOVA; with Tukey’s *post hoc* method) was used for comparisons of more than two groups, and the *t*-test (with Welch correction) for comparison of two groups. For comparison of times of peaks of expression, the distribution is often skewed, therefore the Kruskal-Wallis one-way ANOVA (with Dunn’s *post hoc* method) was used for multiple comparisons and the Wilcoxon rank sum test for comparison of two groups. An alpha level of 0.05 was used for all ANOVA tests.

### Luciferase phase plots

To analyze spatial patterns within the organ, we first create space-time intensity plots of the luciferase images before obtaining a phase representation of the plots using a wavelet transform (henceforth called ‘phase plots’). These phase plots allow interpretation of the space-time dynamics of the signal across the length of the organ independent of amplitude fluctuations.

Space-time phase plots of the luciferase data were created as described previously Gould *et al*, 2018, though with some modifications. Most importantly of which, we include a modification that better allows us to section curved roots. The method including modifications is outlined here in its entirety. Unless otherwise specified steps are implemented via custom developed MATLAB (MathWorks, UK) scripts.

### Image pre-processing

A number of image processing steps were applied prior to the extraction of oscillations:

1. Each seedling is cropped into individual image stacks using ImageJ (NIH, USA) in order to facilitate the further analysis.
2. A rectangle ROI encompassing the whole of the organ of interest plus the surrounding background, is defined. When multiple organs are plotted together (Fig 2H, I) the regions are defined so that there is neither longitudinal gaps nor overlap between them. The ROI is manually checked for signal from neighboring organs or seedlings. These pixels are removed using ImageJ.
3. A 3-by-3 median filter was applied to images to deal with background intensity spikes supposed to be from cosmic rays and camera sensor imperfections
4. The luminescent signal from the organ is segmented from background pixels by applying a threshold to each image individually. The mean of the intensity count across the whole ROI was used as the threshold value.
5. Small objects remaining in the image that are not connected to the organ are removed by applying a morphological opening algorithm. Connected objects less than 50 pixels are removed.

### Intensity space-time plots

To create the space-time plot, we average the signal across longitudinal sections of the organ. However, because plant organs naturally curve during growth we take our longitudinal sections to be perpendicular to the angle of growth. We do this as follows:

1. For a ROI of dimensions *m,n* (with *m* representing the horizontal dimension and *n* the vertical dimension) the grey-level-weighted centroid across each vertical section (*n*) is calculated as

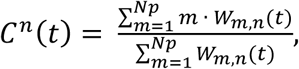

where *W* represents the pixel intensity value and *N*_*p*_ the width of the plant as the number of segmented pixels.
2. A polynomial function of seventh degree is fitted to the centroids to give a curve that describes the shape of the hypocotyl and root {*C*(*t*)} (S2A Fig).
3. At each horizontal position of the ROI {*C*^*n*^(*t*): *n*=1, 2, …)} the tangent and normal line is calculated (S2A Fig).
4. The slope of the normal line is rasterized to give pixel coordinates (S2B Fig). The Bresenham algorithm was utilized for this purpose (67), implemented in MATLAB (68).
5. The rasterized line is limited to 10 pixels, centered around the intersect with the root curve fit {*C*(*t*)}. This prevents multiple intersects with the hypocotyl or root.
6. The mean intensity of the pixels corresponding to the coordinates is taken to give the intensity value for section *n* at time *t* in the space-time intensity plots (S2C Fig).

### Phase space-time plots

We use the wavelet transform (69) to obtain phase plots (S2D Fig) from intensity space-time plots (S2C Fig). The continuous wavelet transform is closely related to the Fourier transform. However, unlike the Fourier transform, the continuous wavelet transform does not assume a stationary signal. This allows the observation of more complex signals including non-constant periods. This could be relevant to our data, given that a clocks response to perturbagens may be transient or changing.

Given a time series *V* = (*V*_1_, …, *V*_n_), the continuous wavelet transform of *V* is given by,

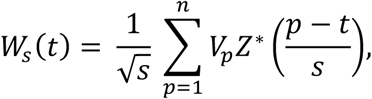

where *Z* is a wave-like function known as the mother wavelet, and *s* is a dimensionless frequency scale variable. *Z*\*denotes the complex conjugate of *Z*. For *Z*, we choose the Morlet wavelet,

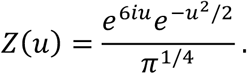

The wavelet transform can instead be expressed in terms of its phase and magnitude,

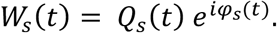

For meaningful interpretation of the phase values *s* must be chosen close to the characteristic period of the times series *V*. However, the resultant phases are robust to small variations of *s*. We therefore select a single *s* for each experimental condition, matching *s* to the frequency of the rhythms that we observe in the root under that condition. Carrying out this procedure for every row of the intensity kymographs results in a phase plot (S2D Fig) corresponding to the intensity plot (S2C Fig). For comparison between plots, we plot the first 16 pixels (approximately 1 mm) of the hypocotyl and the entirety of the root.

### Synchrony analysis

By looking at the all-to-all synchrony between pixels within the hypocotyl and root, the synchrony of oscillators in these tissues can be estimated. We exclude the cotyledons from the analysis because their orientation and movement make phase extraction difficult. For each time point the order parameter (27) *R*, at time *t*, was obtained as

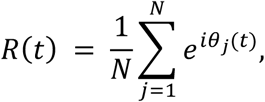

where *N* is the total number of pixels in the hypocotyl and root combined and θ_j_ the phase of the *j*-th pixel. *R* values range from 0 to 1, with a value of 1 indicating a set of completely synchronized oscillators and a value of zero a set of completely desynchronized oscillators.

### Phase oscillator model

As in Gould *et al*., 2018, we use the Kuramoto phase oscillator model to describe the dynamics of *GI*::*LUC* in each pixel (here a pixel represents a set of individual, neighbor cells). We view the plant in 2 dimensions with positions in horizontal and vertical (longitudinal) direction described by index positions *i* and *j*, respectively, so that every pixel, *P*(*i,j*) have an associated position (*i,j*). The phase at the pixel *P*(*i,j*) is represented by *θ*^*(i,j)*^ where its dynamics in time, *t*, are governed by the following equation

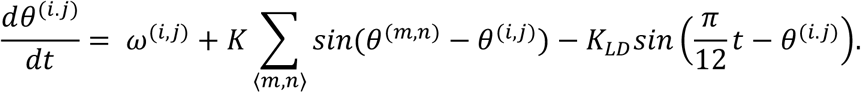

Here the first term is the intrinsic frequency of the pixel, *ω*^*(i,j)*^. The second term is the coupling contribution from the nearest-neighbor pixels in positions (*m,n*) that are closest to (*i,j*), namely, *m*=*i*-1, *i, i*+1 while *n*=*i*-1, *i, i*-1. We assume a plant template that is symmetric and resembles the shape of a seedling (S4 Fig). For sake of simplicity we assume that the coupling constant, *K*, is the same across all pixels and we set it arbitrarily to *K*=1. The final term represents the coupling of the oscillator to the external force, in this case the light force. Here *K*_*LD*_ is the constant for the intensity of the light forcing, where all oscillators are subject to 24 h forcing. Note that when the clocks are not entrained to the LD cycles, *K*_*LD*_=0. Since *GI* tends to peak at onset of dusk in 12 h light-12 h dark cycles and shorter photoperiods (8) we assume that the phase of *GI* will be antiphase to light, hence the negative sign in front of *K*_*LD*_. In our simulations of the LD-to-LD model, we set *K*_*LD*_= 1.

Intrinsic periods are different across different sections of the plant. Intrinsic periods of the pixels in each section are taken from Normal distributions with means of 23.82 h, 25.41 h, 29.04 h and 26.90 h for cotyledon, hypocotyl, root and root tip pixels respectively, with standard deviation at 10 % of the mean value, respectively. The root tip is 5 pixels long and wide.

Initial values of all phases in the LD-to-LL and LD-to-LD simulations are at the time of the start of measurement identical, with first peaks occurring approximately 11 h after the first measurement. In the LL-to-LL model, since we have no information about the phases, we set them to be uniformly distributed across a cycle (i.e. random). We note that in the LL-to-LL model, setting the phases to be in phase, or close to in phase (e.g. approximately 11 h after first measurement ± 2 h (standard deviation)), we could not obtain the results seen. ODEs are solved using the Euler method and simulations were performed in MATLAB.

Since the seedlings in our experiments grow, here we also introduce growth to the template seedling: we allow the root to grow by 1 pixel every five hours. Every newborn cell (and hence the new pixel) has the same phase as the closest set of cells (pixels) in the template, namely new pixels *P*(*i,j*), *P*(*i*+1, *j*), *P*(*i*+2, *j*) will inherit the phases from *P*(*i,j*-1), *P*(*i*+1, *j*-1) and *P*(*i*+2, *j*-1), respectively. Their periods will be taken from the Normal distribution with the mean 26.90 h and the standard deviation of 10 % of the mean value.

After root growth, the root tip should stay fixed in size (of 5-by-5 pixels), so the previous most upper set of root tip pixels at the root/root tip junction will from now on be considered as root tissue instead. This means that their periods lengthen and they will be chosen from a Normal distribution with the mean of 28.04 h and the standard deviation of 10 % of the mean value.

The expression of *GI* for each pixel, *GI*^*(i,j)*^,is calculated from the phase model as: *GI*^*(i,j)*^(*t*) = *cos*(θ^(*i,j)*^ (*t*)) + 1. It follows that the total sum of the luminescence for every longitudinal position *j* is 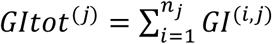 where total number of cells measuring across that section of the plant is *n*_*j*_. The total luminescence is normalized so the maximum peak of expression in every longitudinal position is 1. The phases are extracted from the luminescence using the wavelet transform, as described above for the experimental data in *Phase space-time plots*.

To calculate the periods of the tissues as shown in Fig 4A, B we take regions of 5- by-5 pixels in each tissue (S4C Fig) and calculate the median *GI* expression level for each region. Periods are calculated as the mean of the peak-to-peak periods of the median trace.

An alternative model which could give rise to the LD-to-LL spatial wave behaviors observed is one where there is no coupling but periods increase towards the middle of the root. This means that *K*=0, and we set periods in the root to increase linearly from 25.41 h at the hypocotyl/root junction to 28.04 h in the middle of the root, and then decrease linearly again to 26.90 h at the root/root tip junction. All other previous assumptions are adopted. Here, though a bow-shaped wave of expression can be obtained in the LD-to-LL simulations (S5C and S5D Fig), the model fails to reproduce the behavior observed in LL-to-LL (S5E and S5F Fig).

## Acknowledgments

We thank Alex A. R. Webb (University of Cambridge) and Ozgur E. Akman (University of Exeter) for critical reading of the manuscript and Laszlo Kozma-Bognar (Hungarian Academy of Sciences) for gifts of transgenic material.

## Data and code availability

Project code and datasets will be available from the project GitLab page (www.gitlab.com/slcu/teamJL/greenwood_etal_2019). The following MATLAB File Exchange submissions were also used for the making of figures: ‘shadedErrorBar’ (70), ‘legendflex’ (71), and ‘Alternative box plot’ (72). Fig 1 utilized graphics available from the Plant Illustrations repository (73).

## Abbreviations

LD: light-dark
GI: GIGANTEA
LUC: LUCIFERASE
LL: constant light
h: hours
DD: constant darkness
DCMU: 3-(3,4-dichlorophenyl)-1,1-dimethylurea
MS: Murashige & Skoog
ROI: region of interest
FFT-NLLS: Fast-Fourier Transformed Non-Linear Least Squares
RLL: constant red light
BLL: constant blue light.

---

**Fig S1.**
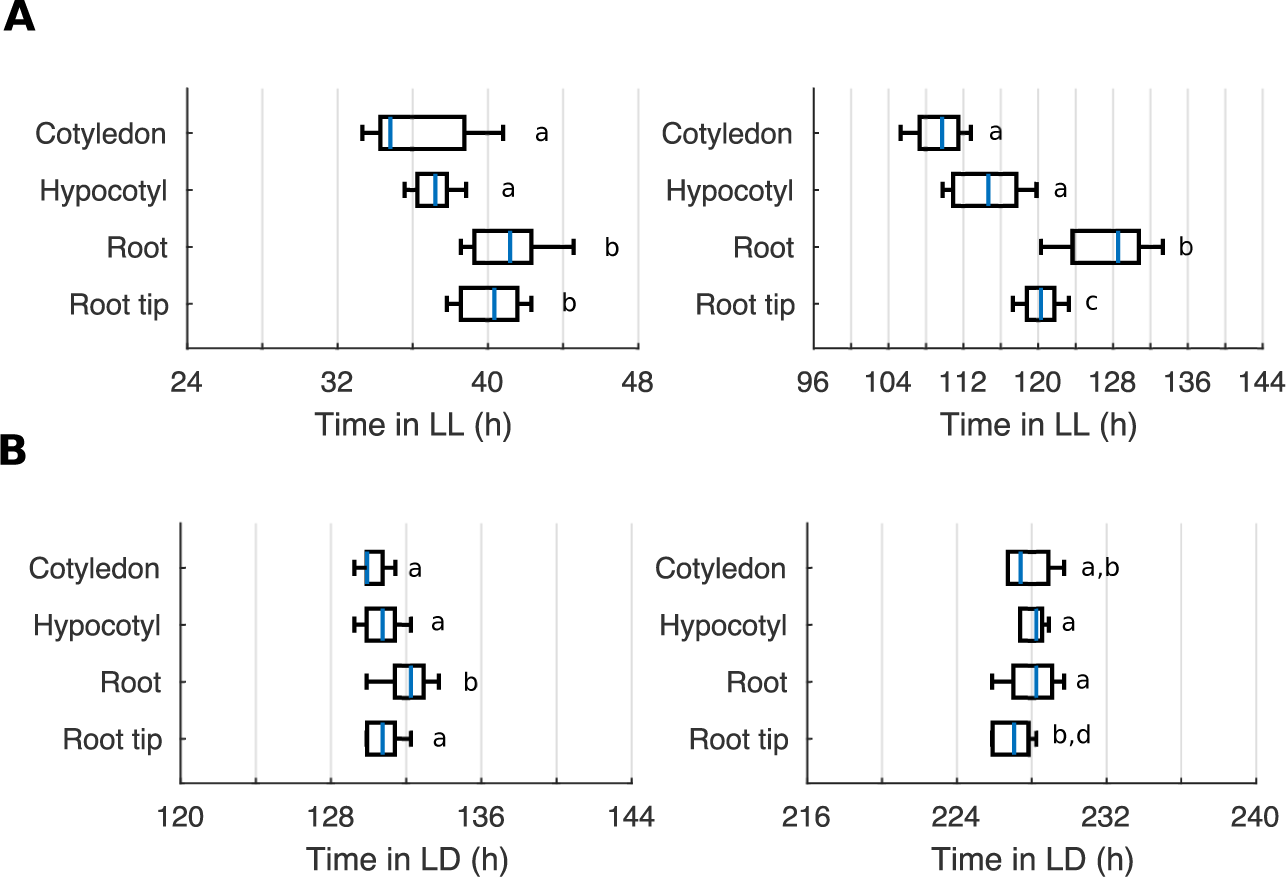
Organ specific clocks show phase differences under constant environmental conditions and light-dark cycles from the first to the final oscillation. A. Times of peaks of expression in different organs during the first (left) and final (right) observed oscillation under LD-to-LL condition. Means are statistically different (*p* < 0.05, one-way ANOVA, Tukey’s *post ho*c tests) if they do not have a letter in common. B. Times of peaks of expression in different organs during the first (left) and final (right) observed oscillation under LD-to-LD condition. Means are statistically different (*p* < 0.05, one-way ANOVA, Tukey’s *post ho*c tests) if they do not have a letter in common. See S1 and S2 File for exact *n* and test statistics. All boxplots indicate the median, upper and lower quartile, and whiskers the 9^th^ and 91^st^ percentile.

**Fig S2.**
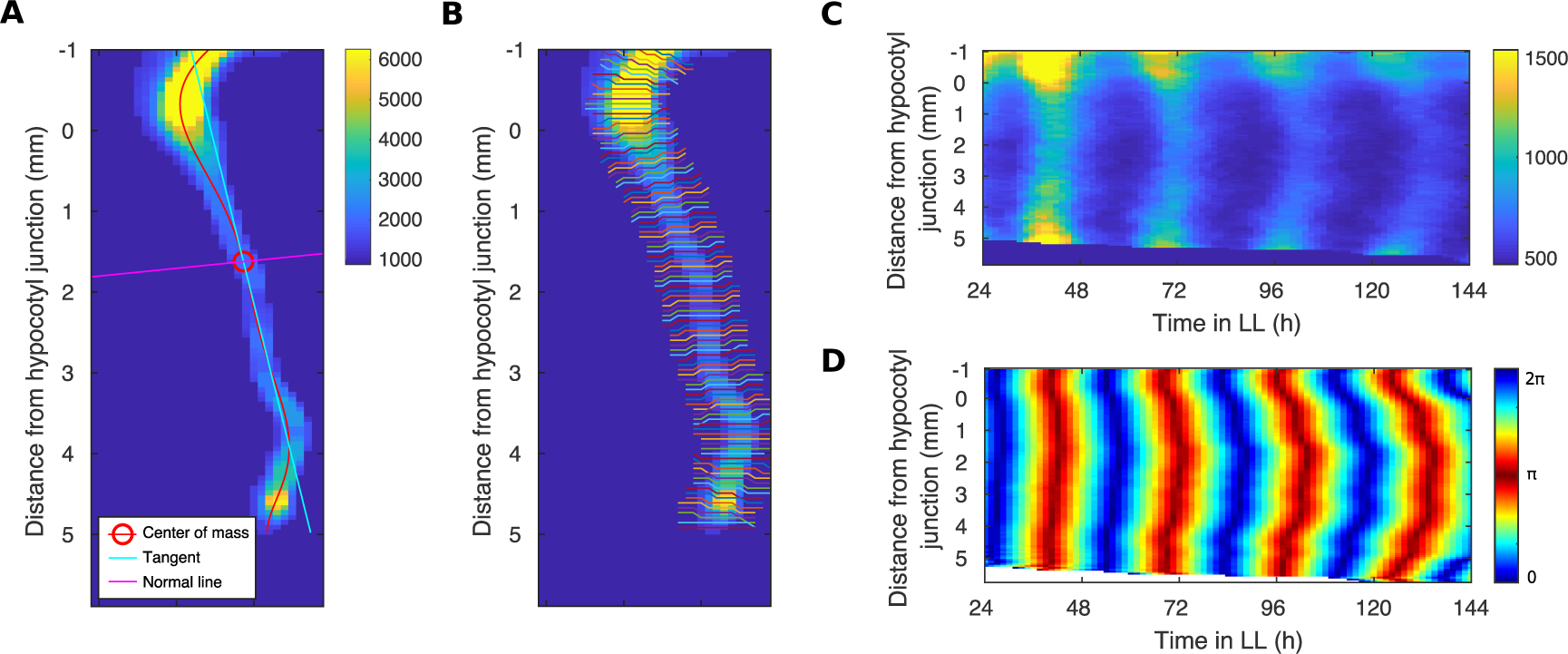
Space-time phase plots from luciferase images. A. Luciferase images are thresholded and a line fitted through the center of mass of the tissue. At each index on this line, the normal line is taken. B. Each normal line is rasterized and limited to 5 pixels around the center of mass to give pixel coordinates for longitudinal sections. C. The mean value across longitudinal sections is taken at each time point to create a raw intensity space-time plot of a single seedling. D. The phase of the oscillations is extracted using a wavelet transform to give a space-time map of the phase.

**Fig S3.**
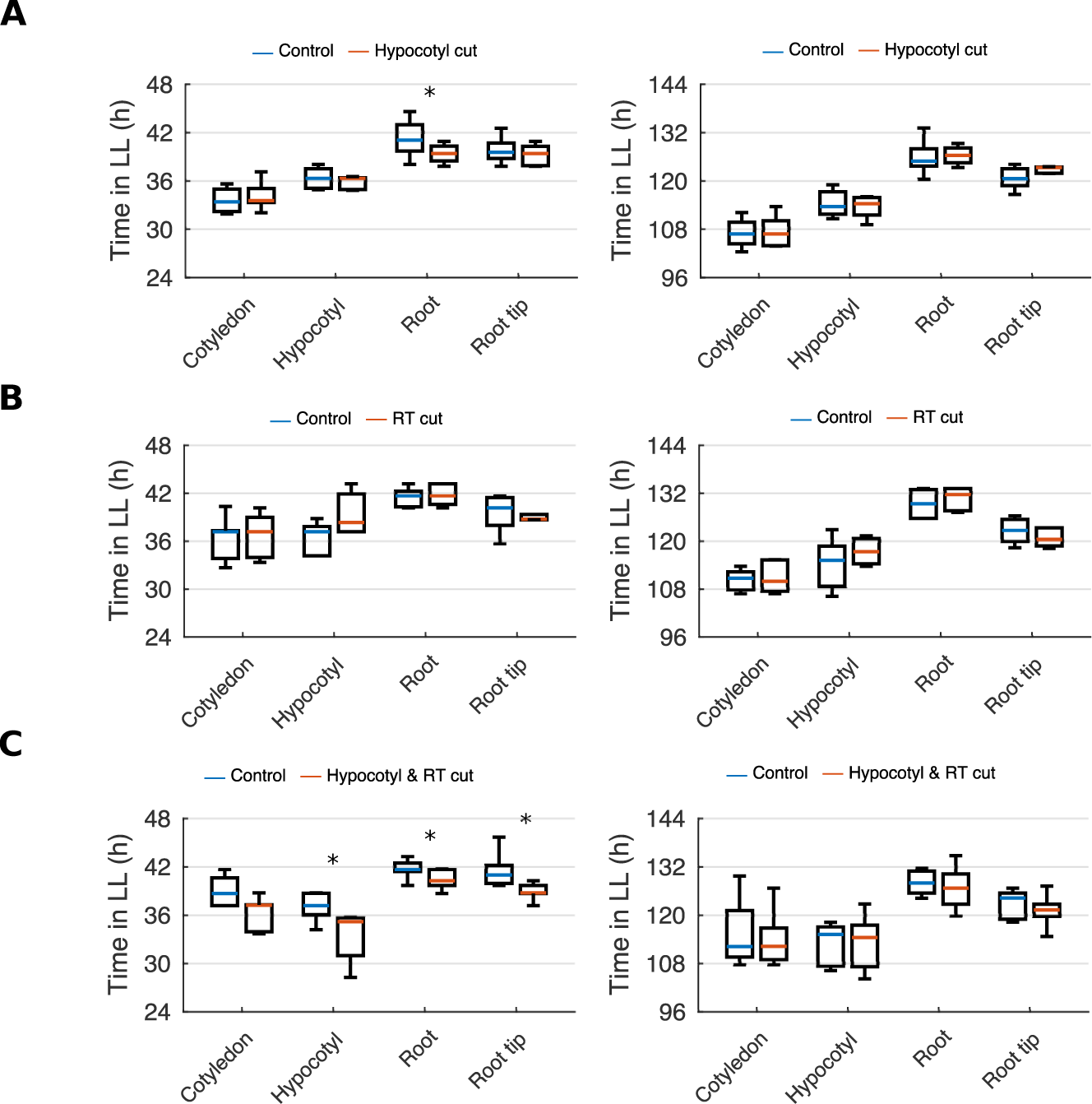
Phase differences between organs following cuts is comparable to controls. A-C. Times of peaks of expression in different organs for the first (left) and final (right) observed oscillation following a cut at the hypocotyl junction (A), root tip (B), or both the hypocotyl junction and root tip (C) conditions. * *p* < 0.05, Wilcoxon rank-sum test. See S1 and S2 File for exact *n* and test statistics. All boxplots indicate the median, upper and lower quartile, and whiskers the 9^th^ and 91^st^ percentile.

**Fig S4.**
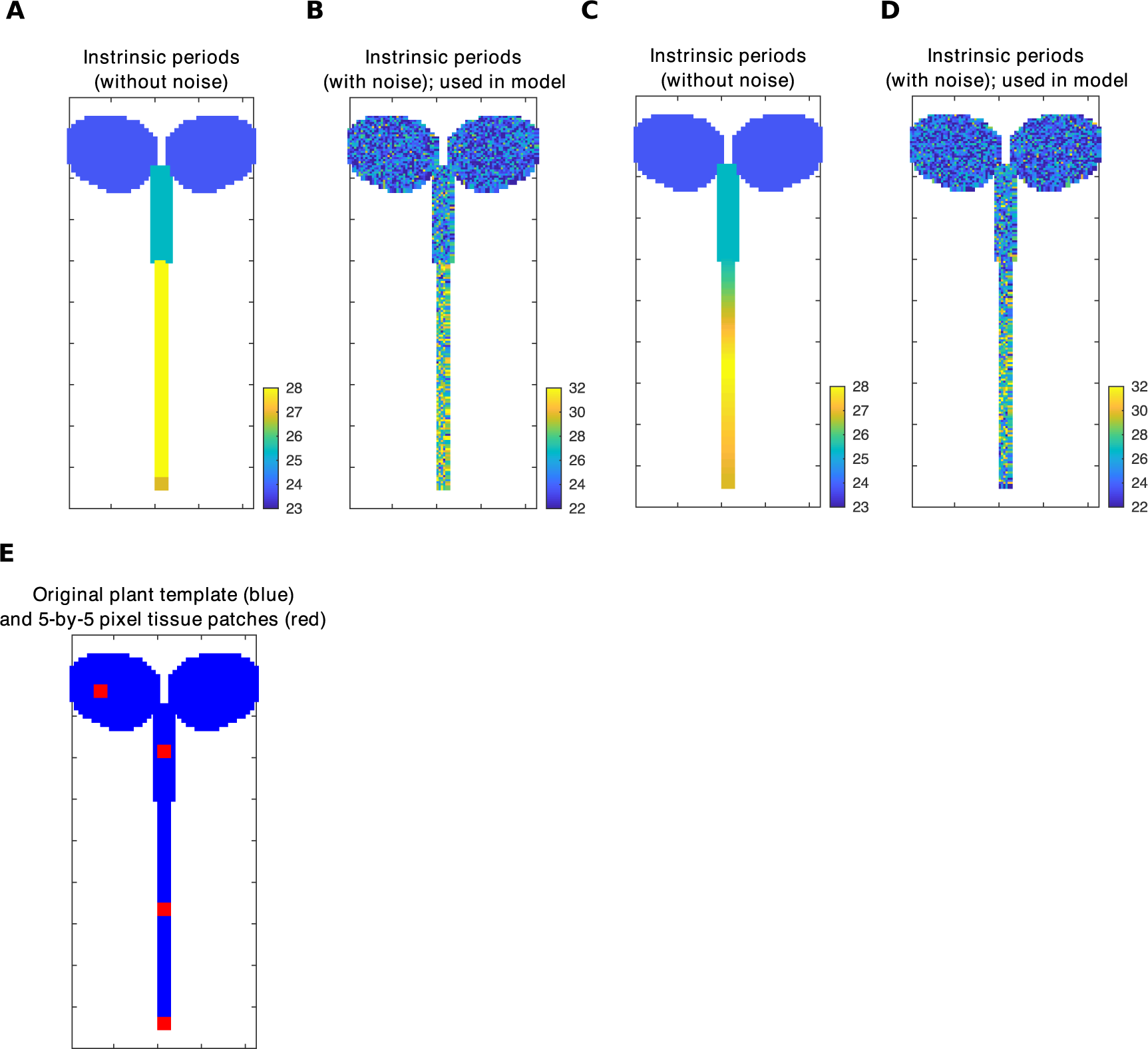
Template for simulations with organ specific periods and the ROI used for analysis. A, B. Template for simulations where in (A) periods of the pixels in each tissue are set to the mean periods measured in the LD-to-LL experimental data. In (B), a representative set of periods for each region are shown, as drawn from the period distributions described in Materials and Methods. C, D. Template for simulations of the alternative model where in (C) periods of the pixels in each tissue are set to the mean periods measured in the LD-to-LL experimental data, but with a gradient of periods in the root as described in Materials and Methods. In (D), a representative set of seedling periods are shown, drawn from the period distributions and gradient described in Materials and Methods. E. The 5-by-5 pixel ROIs used for phase and period analysis are identified on the template.

**Fig S5.**
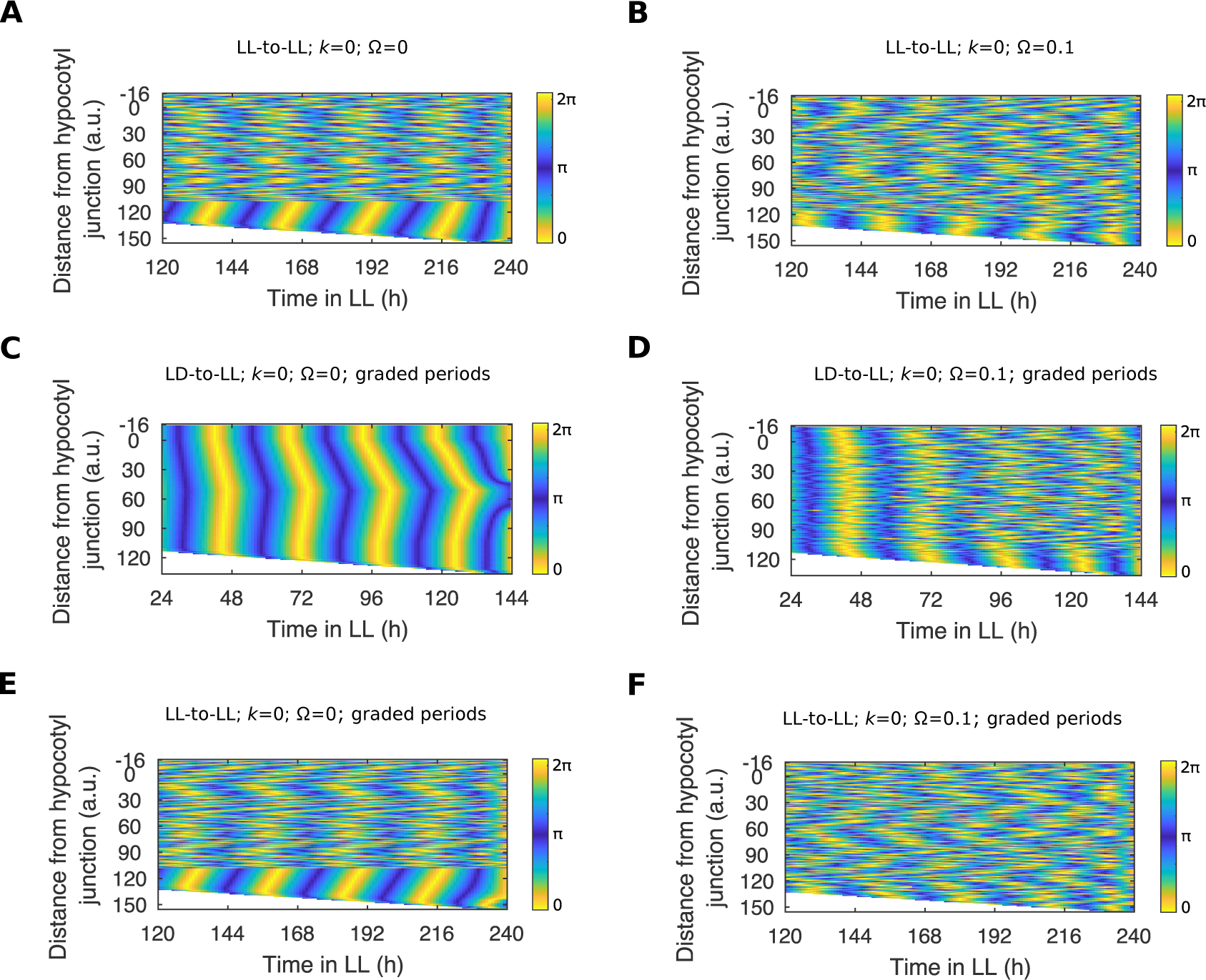
Alternative model simulations. A, B. Phase plot of simulated *GI* expression across longitudinal sections of the hypocotyl and root of a single seedling for LL-to-LL condition in the absence of coupling (*k* = 0), but with period differences. In A, periods of the pixels in each tissue are set to the mean periods measured in the LD-to-LL experimental data, without noise (Ω = 0). In B, a representative set of periods for each region are shown, as drawn from the period distributions described in Materials and Methods (Ω = 0.1). C, D. Phase plot of simulated *GI* expression across longitudinal sections of the hypocotyl and root of a single seedling for LD-to-LL condition in the absence of coupling (*k* = 0). In C, periods in the root region are graded with a maximum period in the middle of the root, without noise (Ω = 0). In D, periods are also graded in the root but periods are drawn from a distribution (Ω = 0.1). See Materials and Methods for details. E, F. Phase plot of simulated *GI* expression across longitudinal sections of the hypocotyl and root of a single seedling for LL-to-LL condition in the absence of coupling (*k* = 0). In E, periods in the root region are graded with a maximum period in the middle of the root, without noise (Ω = 0). In F, periods are also graded in the root but periods are drawn from a distribution (Ω = 0.1). See Materials and Methods for details.

**Fig S6.**
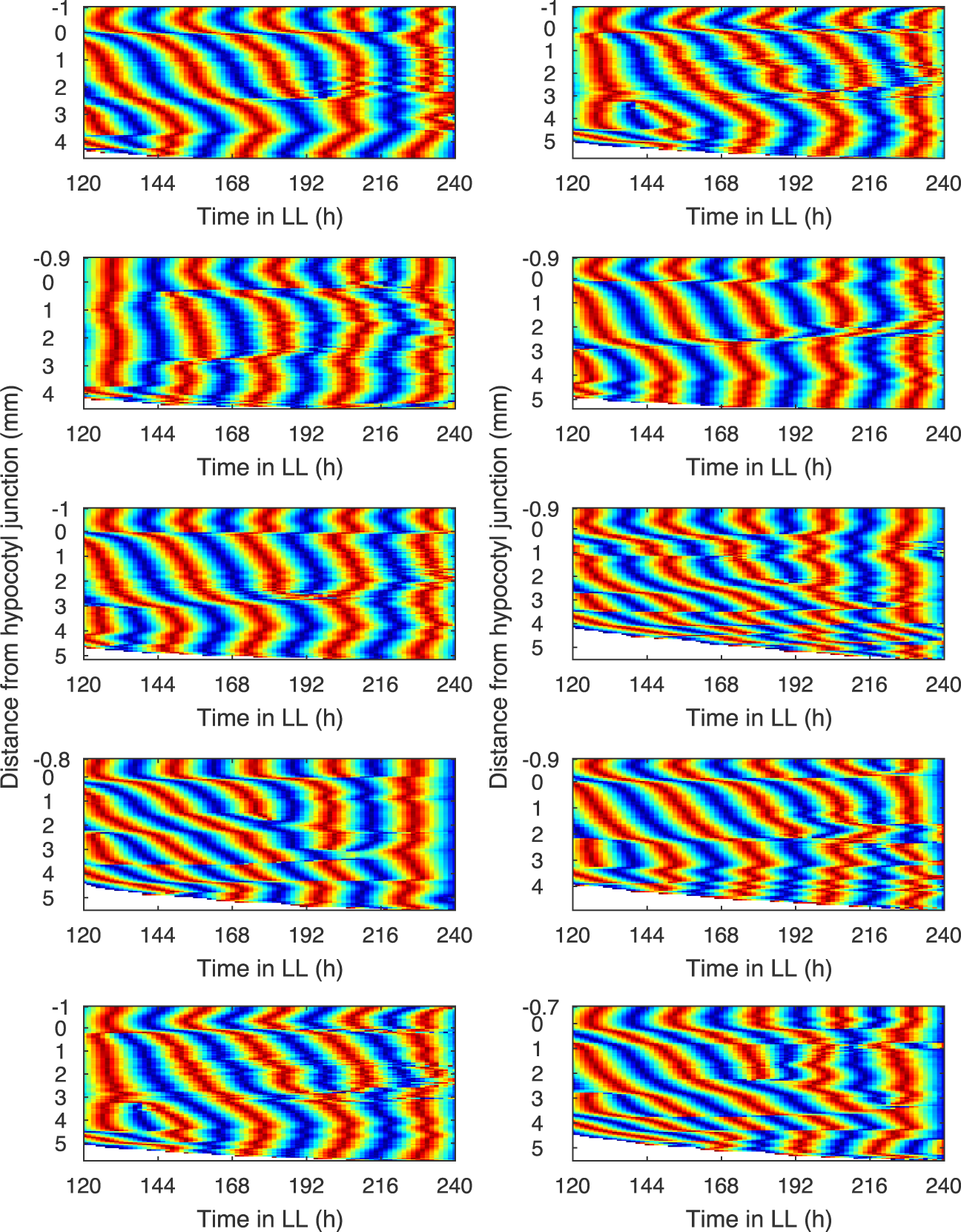
Representative phase plots for LL-to-LL condition. Phase plots of *GI::LUC* expression across longitudinal sections of the hypocotyl and root. Each phase plot is of a single seedling that is representative for the LL-to-LL condition.

**Fig S7.**
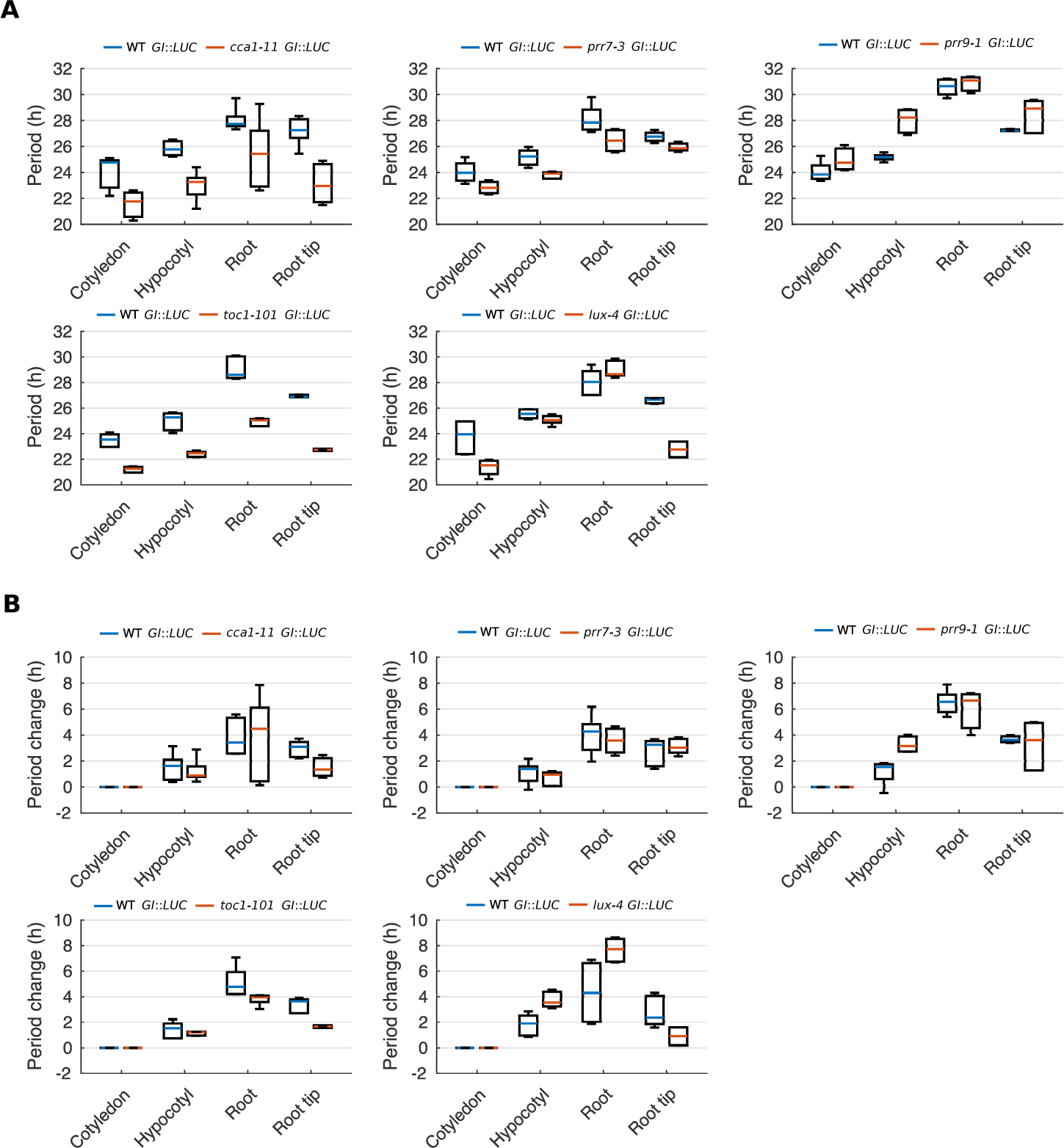
Core clock network mutations affect the period of different regions proportionately. A. Period estimates for *GI*::*LUC* expression from different organs imaged under LD-to-LL condition in circadian mutants lines. B. Period change relative to the cotyledon for *GI*::*LUC* expression from different organs imaged under LD-to-LL condition in circadian mutant lines. For *cca1-11, N* = 4; *prr7-3, N* = 4; *prr9-1, N* = 2; *toc1-101, N* = 2; *lux-4, N* = 2. For all, *n* ≈ 12. *N* represents the number of independent experiments, *n* the total number of seedlings. See S1 and S2 File for exact *n* and test statistics. All boxplots indicate the median, upper and lower quartile, and whiskers the 9^th^ and 91^st^ percentile.

**Fig S8.**
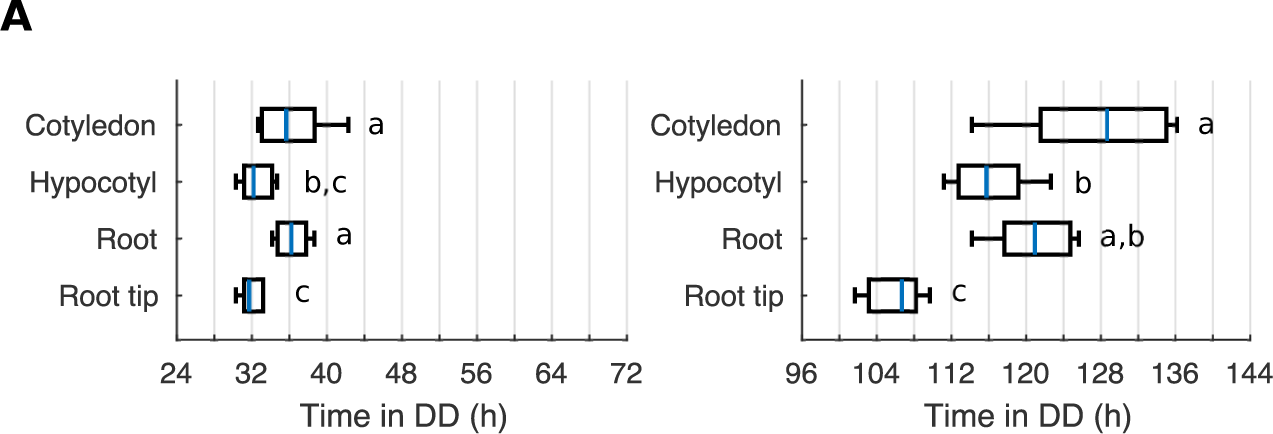
Phase shifts between aerial organs and the root are reduced under constant darkness. A. Times of peaks of expression in different organs during the first (left) and final (right) observed oscillation under constant darkness. Means are statistically different (*p* < 0.05, one-way ANOVA, Tukey’s *post ho*c tests) if they do not have a letter in common. See S1 and S2 File for exact *n* and test statistics. Boxplots indicate the median, upper and lower quartile, and whiskers the 9^th^ and 91^st^ percentile.

**Fig S9.**
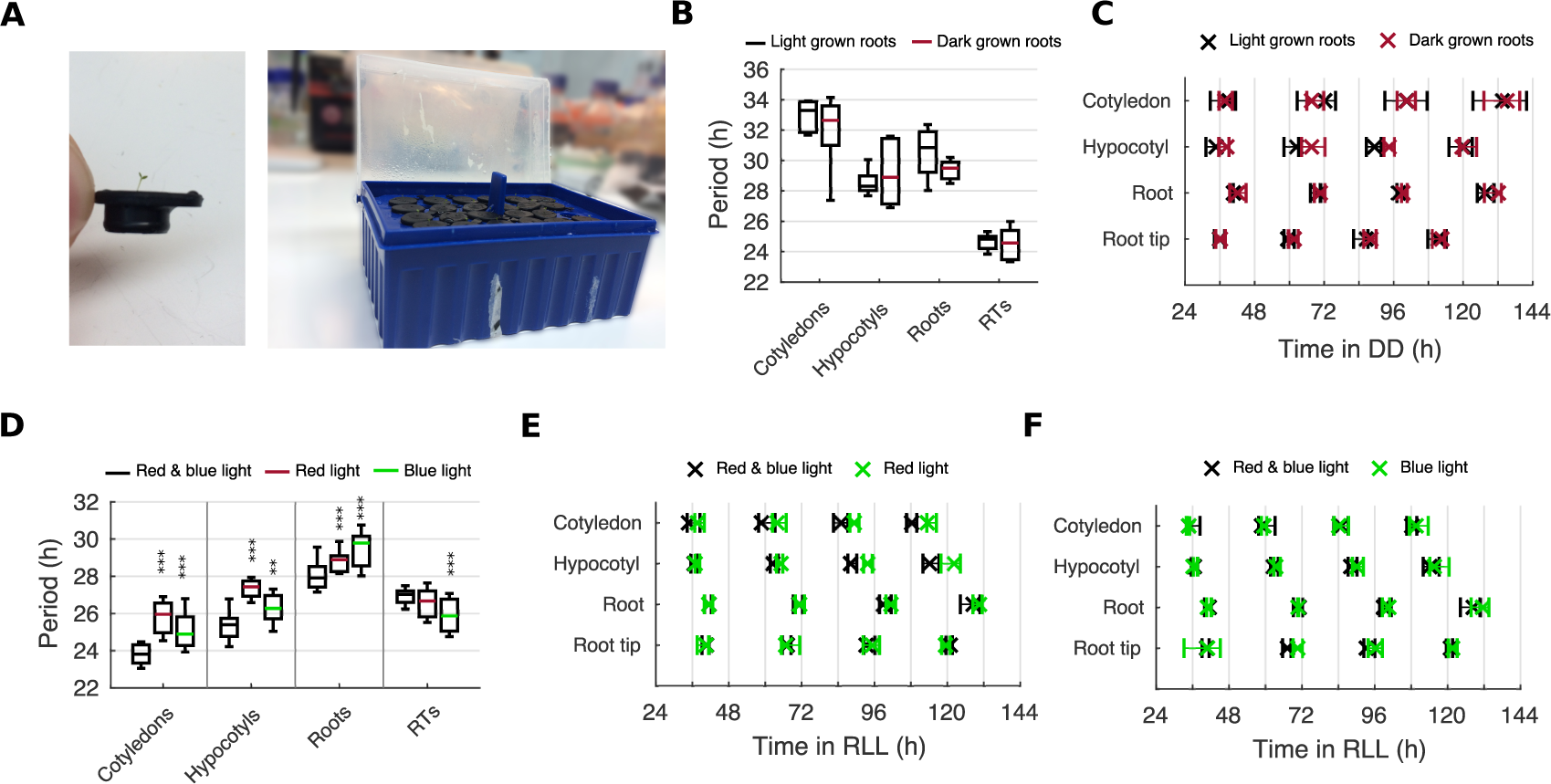
The quality of light input effects rhythms organ specifically. A. Seeds are sown on agar filled black micro-centrifuge tube lids, with a piercing in the lid (left), and suspended in MS liquid in a floating micro-centrifuge tube rack (right), as described previously (62). Note that images include a blur selectively on the background in order to highlight these components. B. Period estimates for different organs in light grown and dark grown roots. All comparisons between period estimates are not significant, *p* < 0.05, by two-tailed *t*-test, Welch correction. C. Times of peaks of expression in different organs for light grown and dark grown roots. Imaging is under constant darkness (DD). Plots represent the 25^th^ percentile, median, and the 75^th^ percentile for the peak times of the oscillations of each tissue. D. Period estimates for different organs under constant red and blue light, red light only, or blue light only. Statistical comparison is to red & blue light data, *** *p* < 0.001, by two-tailed *t*-test, Welch correction. E, F. Times of peaks of expression in different organs imaged under constant red (RLL; B) or constant blue (BLL; C). Plots represent the 25^th^ percentile, median, and the 75^th^ percentile for the peak times of the oscillations of each tissue. For dark grown roots, *N* = 3; RLL, *N* = 2; BLL, *N* = 2. For all, *n* ≈ 20. *N* represents the number of independent experiments, *n* the total number of seedlings. See S1 and S2 File for exact *n* and test statistics. All boxplots indicate the median, upper and lower quartile, and whiskers the 9^th^ and 91^st^ percentile. BLL data is an analysis of time-lapse movies carried out in Gould *et al*., 2018.

**Fig S10.**
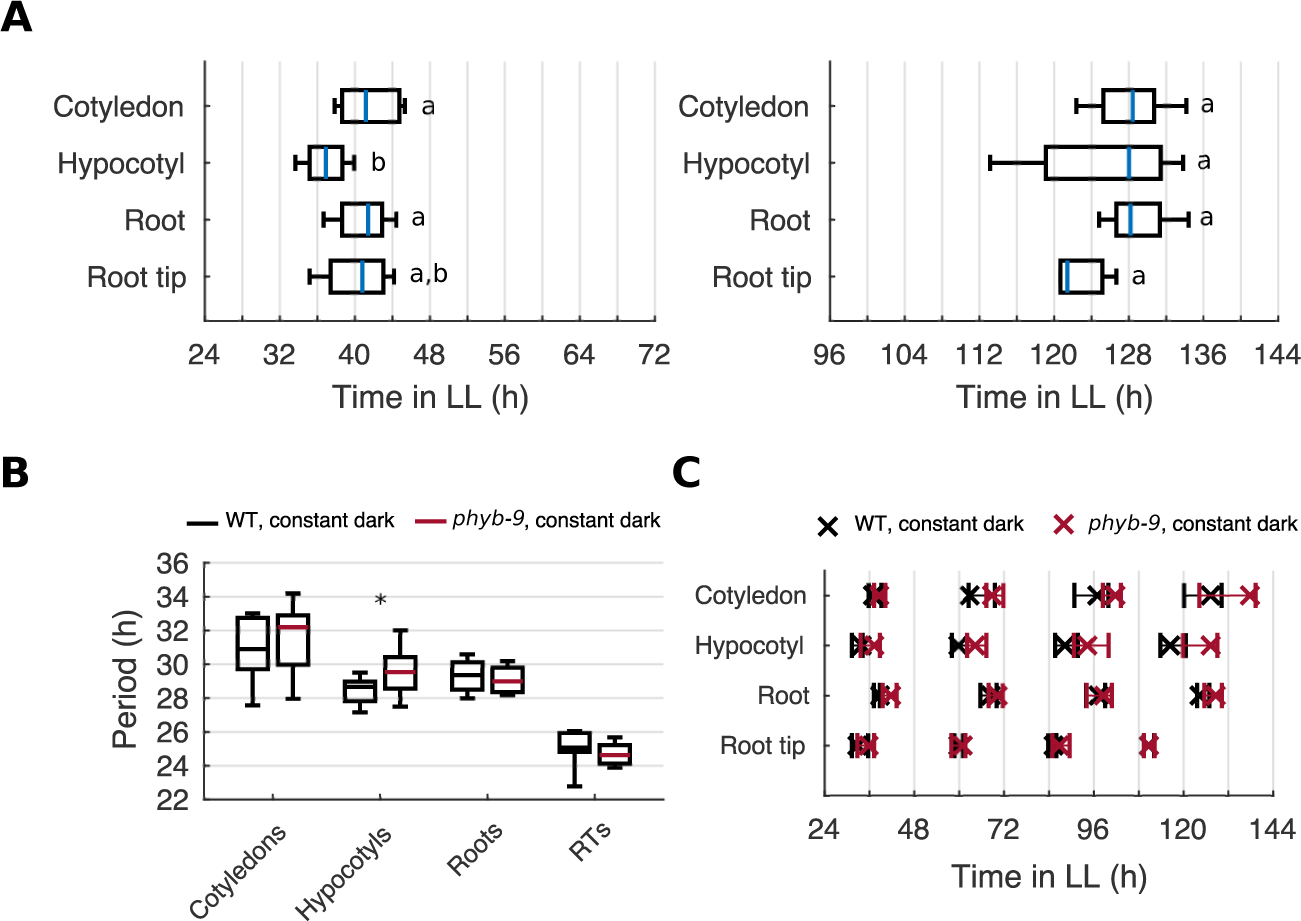
PHYTOCHROME B sets clock periods organ specifically under red light and constant darkness. A. Times of peaks of expression in different organs during the first (left) and final (right) observed oscillation in the *phyb-9* mutant imaged under constant red light. Means are statistically different (*p* < 0.05, one-way ANOVA, Tukey’s *post ho*c tests) if they do not have a letter in common. B. Period estimates for different organs in the *phyb-9* mutant imaged under constant darkness. * *p* < 0.05, by two-tailed *t*-test, Welch correction. C. Times of peaks of expression in different organs in the *phyb-9* mutant imaged under constant darkness. Plots represent the 25^th^ percentile, median, and the 75^th^ percentile for the peak times of the oscillations of each tissue. For *phyb-9* red light, *N* = 4; *phyb-9* DD, *N* = 2; For both *n* ≈ 20. *N* represents the number of independent experiments, *n* the total number of seedlings. See S1 and S2 File for exact *n* and test statistics. All boxplots indicate the median, upper and lower quartile, and whiskers the 9^th^ and 91^st^ percentile.

**Fig S11.**
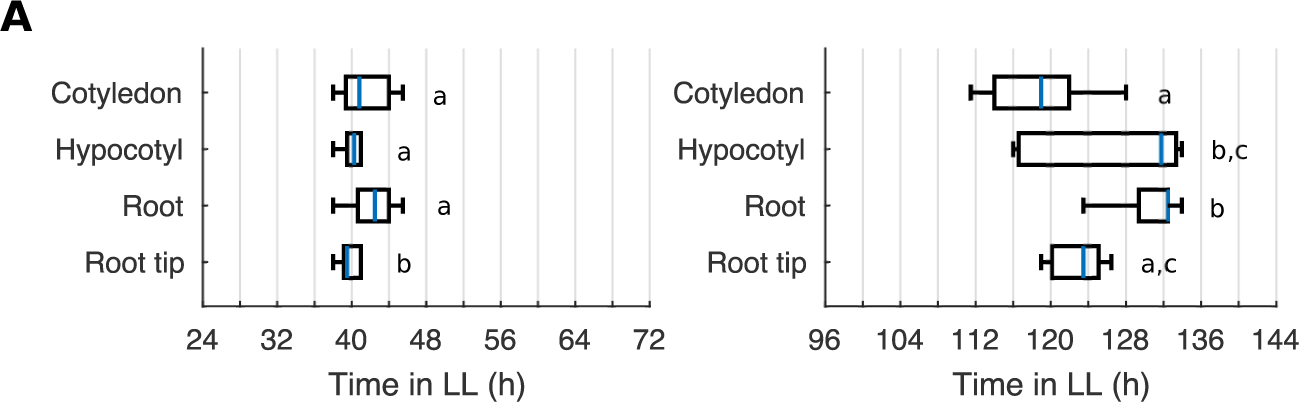
Phase shifts between aerial organs and the root are reduced following the inhibition of photosynthesis by DCMU. A. Times of peaks of expression in different organs during the first (left) and final (right) observed oscillation during the inhibition of photosynthesis by DCMU. Means are statistically different (*p* < 0.05, one-way ANOVA, Tukey’s *post ho*c tests) if they do not have a letter in common. See S1 and S2 File for exact *n* and test statistics. Boxplots indicate the median, upper and lower quartile, and whiskers the 9^th^ and 91^st^ percentile.

**Fig S12.**
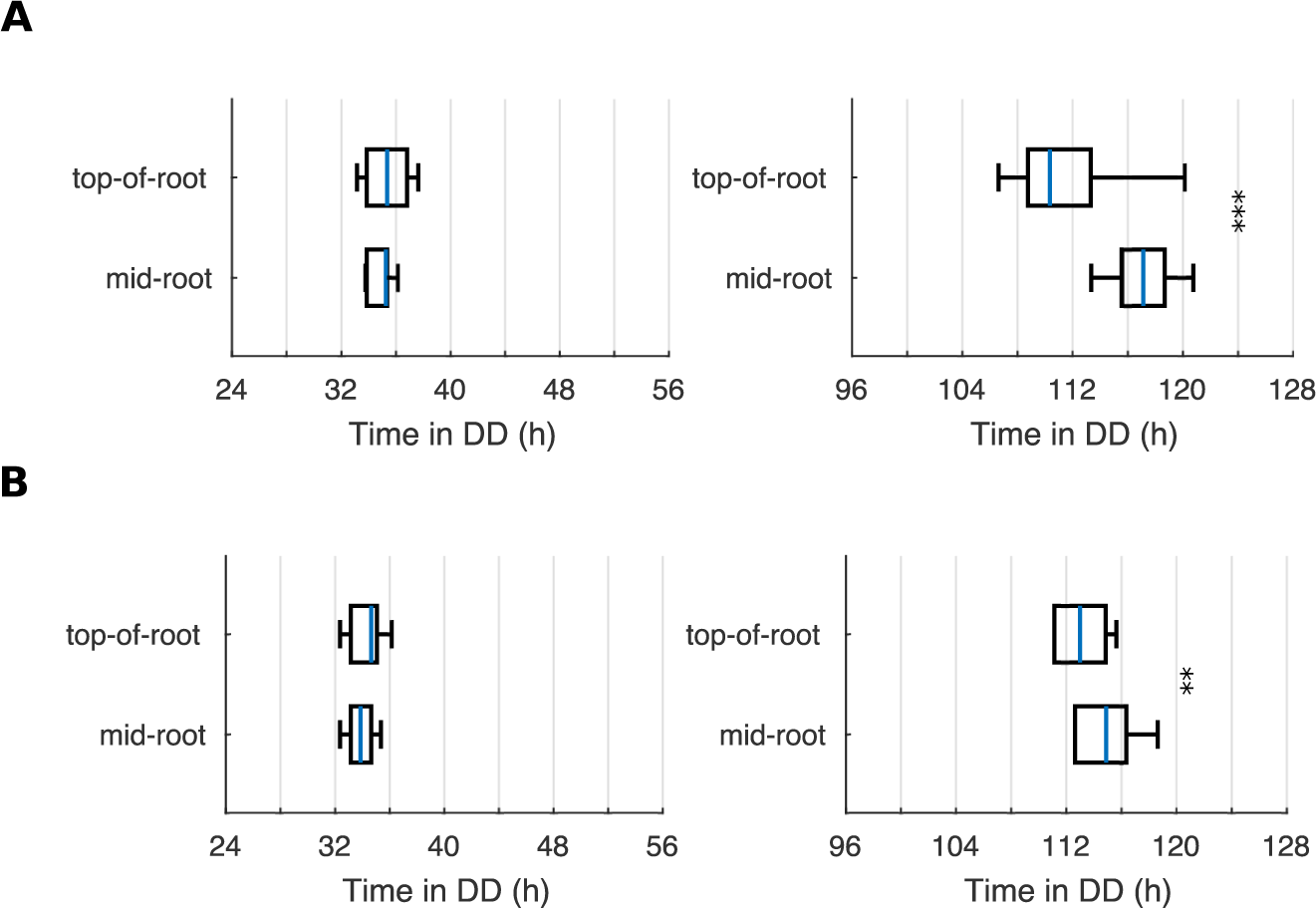
The application of sugar to the top of the root creates a phase shift from the top to the middle of the root under constant darkness. A. Times of peaks of expression in different regions during the first (left) and final (right) observed oscillation during the partial contact of the root with sucrose. *** *p* < 0.001, Wilcoxon rank-sum test. B. Times of peaks of expression in different regions during the first (left) and final (right) observed oscillation during the partial contact of the root with mannitol. ** *p* < 0.01, Wilcoxon rank-sum test. See S1 and S2 File for exact *n* and test statistics. Boxplots indicate the median, upper and lower quartile, and whiskers the 9^th^ and 91^st^ percentile.

**S1 Video. Spatial waves of *GI*::*LUC* expression under the LD-to-LL condition.** *GI*::*LUC* luminescence from 24–144 h after transfer to constant light. Frame intervals are 90 minutes and scale bar shows 0.5 mm.

**S2 Video. Spatial waves of *GI*::*LUC* expression under the LD-to-LD condition.** *GI*::*LUC* luminescence from 24–144 h after transfer to constant light. Frame intervals are 90 minutes and scale bar shows 0.5 mm.

**S3 Video. Spatial waves of *GI*::*LUC* expression in a cut root.** *GI*::*LUC* luminescence from 24–144 h after transfer to constant light, following following excision of the root tip 2 h after transfer to constant light. Frame intervals are 90 minutes and scale bar shows 0.5 mm.

**S4 Video. Spatial waves of *GI*::*LUC* expression under the LL-to-LL condition.** *GI*::*LUC* luminescence from 24–144 h after transfer to constant light. Frame intervals are 90 minutes and scale bar shows 0.5 mm.

**S5 Video. Spatial waves of *GI*::*LUC* expression under constant darkness.** *GI*::*LUC* luminescence from 24–144 h after transfer to constant darkness. Frame intervals are 90 minutes and scale bar shows 0.5 mm.

**S6 Video. Spatial waves of *GI*::*LUC* expression under constant red light in the *phyb-9* background.** *GI*::*LUC* luminescence from 24–144 h after transfer to constant darkness. Frame intervals are 90 minutes and scale bar shows 0.5 mm.

**S7 Video. Spatial waves of *GI*::*LUC* expression in the root following application of exogenous sucrose to the top of the root.** *GI*::*LUC* luminescence from 24–144 h after transfer to constant darkness. The top portion of the root (approximately 1 mm) is in contact with sucrose supplemented media whilst the remainder of the root is in contact with media without sugar. Frame intervals are 90 minutes and scale bar shows 0.5 mm.

**S8 Video. Spatial waves of *GI*::*LUC* expression in the root following application of exogenous mannitol to the top of the root.** *GI*::*LUC* luminescence from 24–144 h after transfer to constant darkness. The top portion of the root (approximately 1 mm) is in contact with mannitol supplemented media whilst the remainder of the root is in contact with media without sugar. Frame intervals are 90 minutes and scale bar shows 0.5 mm.

**S1 File. Test statistic values relating to period estimates presented in Figures.**

**S2 File. Test statistic values relating to phase estimates presented in Figures.**

